# Pathological axonal enlargement in connection with amyloidosis, lysosome destabilization and hemorrhage is a major defect in Alzheimer’s disease

**DOI:** 10.1101/2024.06.10.598057

**Authors:** Hualin Fu, Jilong Li, Chunlei Zhang, Guo Gao, Qiqi Ge, Xinping Guan, Daxiang Cui

## Abstract

Alzheimer’s disease (AD) is a multi-amyloidosis disease with Aβ deposits in the cerebral blood vessels, microaneurysms and senile plaques. How Aβ amyloidosis affects axon pathology is not well-examined. We studied Aβ-related axonal phenotypes with histochemistry, immunohistochemistry and fluorescence imaging methods. Widespread axonal amyloidosis with distinctive axonal enlargement was observed in AD. Aβ-positive axon diameters in AD brains were 1.72 times of control brain axons by average. Axonal amyloidosis also associated with MAP2 reduction, Tau phosphorylation, lysosome destabilization and hemorrhagic markers such as ApoE, HBA, HbA1C and Hemin. Lysosome destabilization in AD was also clearly identified in the neural soma, associating with the co-expression of Aβ and Cathepsin D, HBA, ACTA2 and ColIV, implicating exogeneous hemorrhagic protein intake might influence neural lysosome stability. The data showed that Aβ-containing lysosomes were 2.23 times as large as the control lysosomes. Furthermore, under rare conditions, axon breakages were observed, which likely resulted in Wallerian degeneration. In summary, axonal enlargement associated with amyloidosis, chronic microhemorrhage and lysosome destabilization is a major defect in Alzheimer’s disease.

## Introduction

The pathological hallmarks of Alzheimer’s disease include senile plaques, neurofibrillary tangles and neuritc dystrophy[1, 2]. Neuritc dystrophy refers to the dystrophy of axons and dendrites. In our previous investigation, we identified cerebral microaneurysm rupture and hemorrhagic Aβ leakage as a pathological mechanism for senile plaque formation[3]. Yet, how chronic hemorrhagic leakage impacts on other pathological phenotypes such as axonal dystrophy in Alzheimer’s disease is not well-studied. Previous studies implicated that both Aβ and Tau phosphorylation are involved in the axonal defects in Alzheimer’s disease[4–7]. However, the exact linkage between Aβ, Tau and their contribution to axonal dystrophy is still not very clear.

In the current study, we analyzed the pathological phenotypes in axons in AD brain tissues by immunohistochemistry, histochemistry and fluorescent imaging methods. The results showed that the major axonal phenotype in Alzheimer’s disease (AD) is the widespread axonal amyloidosis and enlargement with additional spheroid formation. The axonal defect in AD is associated with the enrichment of Aβ, phos-Tau, destabilized lysosomes and hemorrhage-related proteins in axons.

## MATERIAL AND METHODS

### Tissue sections

AD patient frontal lobe brain paraffin tissue sections were purchased from GeneTex (Irvine, CA, USA). Additionally, AD patient and non-AD control frontal lobe brain paraffin tissue sections were provided by National Human Brain Bank for Development and Function, Chinese Academy of Medical Sciences and Peking Union Medical College, Beijing, China. This study was supported by the Institute of Basic Medical Sciences, Chinese Academy of Medical Sciences, Neuroscience Center, and the China Human Brain Banking Consortium. All procedures involving human subjects are done in accord with the ethical standards of the Committee on Human Experimentation in Shanghai Jiao Tong University and in accord with the Helsinki Declaration of 1975.

### List of antibodies for immunohistochemistry

The following primary antibodies and dilutions have been used in this study: Aβ (Abcam ab201061, 1 : 200, Cambridge, UK), Aβ/AβPP (CST #2450, 1 : 200, Danvers, MA, USA), phos-Tau (Abcam ab151559, 1:200), MAP2 (Proteintech 17490-1-AP, 1:200, Rosemont, IL, USA), ApoE (Abcam ab183597, 1:200), Sortilin1 (Abcam ab263864, 1:200), HBA (Abcam ab92492, 1:200), Cathepsin D (Abcam ab75852, 1:200), Lamp2 (Proteintech, 66301-1-Ig, 1:100), Cathepsin D (Proteintech 66534-1-Ig, 1 : 100), HbA1c (OkayBio K5a2, 1 : 100, Nanjing, Jiangsu, China), Hemin (Absolute Antibody, 1D3, 1 : 100, Redcar, UK), AGE (Advanced Glycation End Products) (Abcam, ab23722, 1:200), ColIV (Abcam ab236640, 1:200), ACTA2 (Proteintech 23081-1-AP, 1:400). The following secondary antibodies and dilutions have been used in this study: donkey anti-mouse Alexa-594 secondary antibody (Jackson ImmunoResearch 715-585-150, 1:400, West Grove, PA, USA), donkey anti-rabbit Alexa-488 secondary antibody (Jackson ImmunoResearch 711-545-152, 1:400), donkey anti-rabbit Alexa-594 secondary antibody (Jackson ImmunoResearch 711-585-152, 1:400), and donkey anti-mouse Alexa-488 secondary antibody (Jackson ImmunoResearch 715-545-150, 1:400).

**Immunohistochemistry** was performed as described[8]. Briefly, paraffin sections were deparaffinized by Xylene, 100% EtOH, 95% EtOH, 75% EtOH, 50% EtOH, and PBS washes. Sections were then treated with 10 mmol/L pH 6.0 sodium citrate or 10 mmol/L pH 9.0 Tris-EDTA antigen retrieval solutions in a microwave at high power for 5 min to reach the boiling point and then at low power for another 15 min. The sections were allowed to naturally cool down to room temperature. Then, the slides were blocked with TBST 3% BSA solution for 1 hour at room temperature. After blocking, the samples were incubated with primary antibodies at room temperature for 2 hrs followed by 5 washes of TBST. After that, the samples were incubated with fluorescent secondary antibodies overnight at 4 degree. The treated samples were washed again with TBST 5 times the second day and mounted with PBS+50% glycerol supplemented with Hoechst nuclear dye (Sigma, B2261, 1 μg/ml) and ready for imaging. IHC experiments without primary antibodies were used as negative controls. All experiments have been repeated in order to verify the reproducibility of the results.

**Rhodanine staining** was performed as described[3]. The Rhodanine staining kit (BA4346) was purchased from Baso Diagnostics Inc. (Zhuhai, China). Briefly, paraffin sections were first deparaffinized as described above. After a quick wash in 95% ethanol, the slides were stained in the Rhodanine staining solution (0.01% p-Dimethylaminobenzalrhodanine) in a 37-degree incubator for 30 minutes. After staining, the slides were treated with a 95% ethanol quick wash and 5 times wash in water, then mounted with PBS+50% glycerol solution for imaging analysis.

### Alizarin Red staining

The Alizarin Red staining kit C0138 was purchased from Beyotime Biotechnology (Shanghai, China). Briefly, paraffin sections were deparaffinized as described above. Then, the slides were stained in the Alizarin Red staining solution (2% Alizarin Red, pH4.2) for 5 minutes at room temperature. After staining, the sections were washed 5 times with water, then mounted with PBS+50% glycerol solution for imaging.

### Imaging and morphometry analysis

The fluorescent images were captured with a CQ1 confocal fluorescent microscope (Yokogawa, Ishikawa, Japan) and the color images of sections with histological stains were taken with an Olympus IX71 fluorescent microscope (Olympus Co., Toyoko, Japan). There is a difference in terms of exposure settings when imaging axonal staining versus senile plaque staining. To image axonal Aβ staining clearly, the exposure time should be increased significantly. Images were then analyzed with ImageJ software (NIH, USA). When comparing signal intensities with the same exposure settings, the mean area density was used as the parameter to define the marker densities. The perimeters of axons in transverse orientations were measured to simplify the axon diameter estimation by using the following formula: diameter = perimeter/3.14. The diameters of axons could also be measured directly with the ImageJ line tool utilizing the microscopy scale as a reference.

### Statistics

All data first went through a Shapiro & Wilk normality test using SPSS Statistics 19 software. Two-tailed unpaired T-test was used to compare the means of data with a normal distribution with Excel 2007 software. For data that were not normally distributed, nonparametric Mann-Whitney test was performed to compare the means by using SPSS Statistics 19 software. For the correlation analysis, Pearson correlation was used for the data with a normal distribution while Spearman correlation was used for the data that were not normally distributed. The p Value threshold for statistical significance is set at 0.05. If p<0.05, then the difference was considered statistically significant. When p<0.001, it was labeled as p<0.001. If p≧0.001, then the p Value was shown as it was.

## Funding

This work was supported by the National Natural Science Foundation of China (81472235), the Shanghai Jiao Tong University Medical and Engineering Project (YG2021QN53, YG2017MS71), the International Cooperation Project of National Natural Science Foundation of China (82020108017), and the Innovation Group Project of National Natural Science Foundation of China (81921002).

## Author contributions

HF conceived the study, designed and supervised the experiments and wrote the manuscript; HF, JL performed the experiments; HF, JL, CZ, GG, QG, XG and DC did the data analysis; all authors reviewed the manuscript.

## Competing interests

Authors declare no competing interests.

## Data availability statement

The data that supports the findings of this study are available from the corresponding author upon request.

## Supplementary Materials

Supplementary Figures 1 to 11.

## Results

### Axon amyloidosis and enlargement in AD brain tissues

In our previous publications, Aβ staining was detected in senile plaques, CAA, neurons and red blood cells[3, 9]. After careful re-examination of the Aβ staining patterns in AD brain sections with optimized staining conditions and longer exposure settings, we found that there were clear axon-specific Aβ staining patterns that we previously did not recognize (two overview images of Aβ staining of AD axons in the longitudinal and transverse orientation were displayed in Supplementary Figure 1 and 2). The axon fragments we focused on in this study are mainly axons in the gray matter or at the gray matter/white matter boundary, which could be defined as proximal axons that are close to the neuron soma. The study extended the analysis to axons in the white matter since many aspects of the axon fragments in the white matter showed similar changes as the proximal axons. Figure 1A and Supplementary Figure 1, 2 showed that axon amyloidosis is widespread in AD brain tissues. Aβ immunostaining was distributed along the length of many axons. Aβ-positive axons can be observed both around senile plaques and without senile plaques in the vicinity. Axonal amyloidosis could be detected in axons at both longitudinal and transverse orientation. Axons with Aβ staining were often shown as grouped clusters, corresponding to the known structural character in the human brain cortex as “axon bundles”. Axon bundle, or in the name of “axonal tract” [10] were frequently observed in the Layer VI of the cerebral cortex, which was at the boundary of grey matter/white matter formation. An image of MAP2-labeled axon bundles in the control brain was shown in the Supplementary Figure 3. To our knowledge from the literature, “axon bundles” derived from the neurons is the only, type of neural cell processes that shows a “bundling” phenotype, different from astrocyte or microglial processes (Supplementary Figure 4). These intracortical axon bundles had an average transverse diameter of 61.61±9.88 um (N=11). Each bundle contained an average of 19.09±4.61 axons per area (N=11) (Supplementary Figure 3). The enlarged and often bundled axons but not unaffected axons in AD brain tissues could also be clearly identified with phase contrast microscopy because they have enhanced contrast and are darker in appearance. These enlarged axons were not labeled with microglial marker Iba1 or astrocyte marker GFAP (Supplementary Figure 4). Additionally, the enlarged axons could be identified with an alternative marker of amyloidosis other than Aβ antibody staining, the amyloid blue autofluorescence[11, 12], which we referred to a term of “MetaBlue” in the previous studies[3, 9]. From cortical surface to white matter, axon bundles at transverse orientation or longitudinal orientation could be observed within a same pathological section indicating different populations of axon bundles (Supplementary Figure 1, 2, 3). Axon bundles were most clearly observed in Layer VI of the brain cortex, showing as long axon fragments that bundled together.

**Figure 1:**
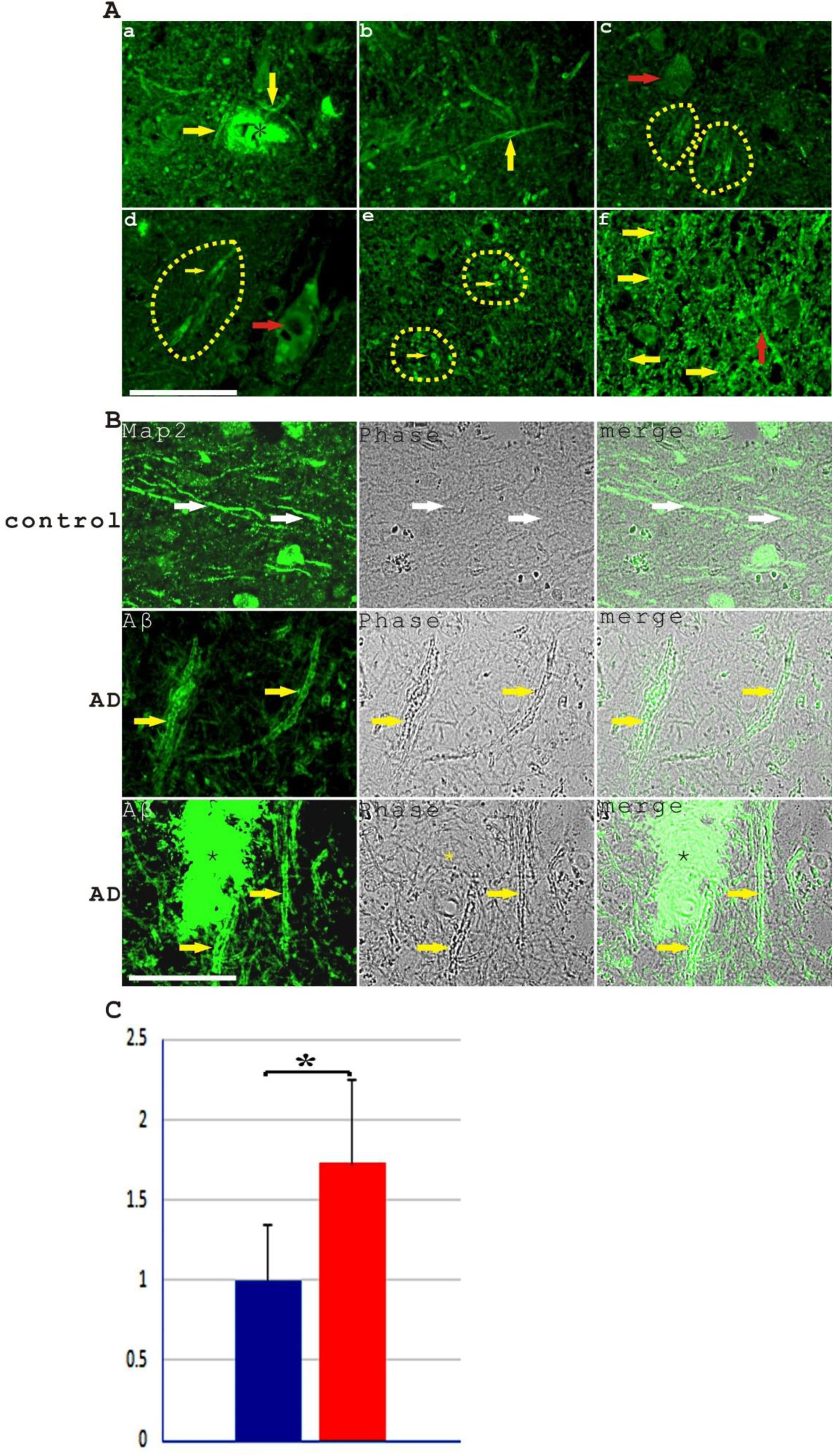
Axon amyloidosis and enlargement are important pathological phenotypes in Alzheimer’s disease. **A.** Axon amyloidosis was detected in AD frontal brain tissues with Aβ antibody with green fluorescence. **a**, Aβ expression was detected both in a senile plaque (indicated with an asterisk) and in axons (indicated with arrows). **b**, An axon at the longitudinal orientation showed spheroid formation stained with Aβ antibody (indicated with an arrow). **c**, Two clusters of Aβ-positive axons in the longitudinal direction (indicated by dashed lines) were shown, indicating Aβ staining of axon bundles. An Aβ-containing neural cell was indicated with a red arrow. **d**, A cluster of Aβ-positive axons in the longitudinal direction (indicated by dashed lines) was shown while blood vessel lumen Aβ expression was indicated with a red arrow. **e**, Two clusters of Aβ-positive axons in the transverse direction (indicated by dashed lines) were shown while individual axons were indicated by yellow arrows. **f,** Axons with abundant axonal Aβ staining mostly in the transverse direction (yellow arrows) were observed in the white matter with one longitudinal axon indicated by a red arrow. Scale bar, 50 μm. **B**. A general enlargement of axons in AD brain tissues was observed when comparing control axons (indicated by white arrows, top panel) marked by MAP2 antibody staining to amyloid-loaded axons in AD (indicated by yellow arrows, bottom two panels) marked by Aβ immunostaining. A senile plaque in the bottom panel was labeled with an asterisk. Scale bar, 50 μm. **C**. Imaging measurements showed that Aβ-positive axons had an average diameter of 1.94±0.59 μm (N=200) while MAP2-labeled control axons measured around 1.13±0.39 μm (N=34). Aβ-positive axons were significantly enlarged, averaging 1.72±0.52 fold of control axons (p<0.001, Mann-Whitney test).

Moreover, puff-like spheroid formation can be detected in a subset of the Aβ-positive axons. Spheroids are often formed at positions with further-enhanced Aβ staining. Aβ-positive axons have a general enlargement phenotype (Figure 1B), which is a basic character appearing more frequently than the spheroid formation. Topologically, “spheroid” could be viewed as an extreme form of “enlargement”. The slight difference of “ enlargement” vs “spheroid” in describing axonal defects could be analogous to the concept of “fusiform” or “saccular form” when describing “aneurysm”. The diameters of Aβ-positive axons in the transverse orientation outside of senile plaque regions were measured since the strong plaque Aβ staining precludes the accurate identification of Aβ-positive axons within the senile plaques. The measured Aβ-positive proximal axons had an average diameter of 1.94±0.59 μm (N=200) while MAP2-labeled proximal axons from control front brain samples measured around 1.13±0.39 μm (N=34). In another word, Aβ-positive axon diameters were around 1.72±0.52 fold of control axons (Mann Whitney test, p<0.001). In addition, among AD axon samples, the mean Aβ staining intensity was weakly positively related to the axon diameter (r=0.338, p<0.001, N=200, Spearman correlation), further emphasizing the positive influence of Aβ on the size enlargement of axons. When axons in the white matter were analyzed, MetaBlue-positive amyloid-laden axon diameters were around 3.80±0.69 μm (N=100), showing even more severe enlargement than the proximal axons in the gray matter or at the gray matter/white matter boundary. The data suggested that the Aβ-amyloidosis-associated axonal enlargement is a basic pathological phenotype in Alzheimer’s disease.

### Axons in AD patients showed a reduction of structural protein MAP2

To examine the effect of axon amyloidosis on the neurons, AD brain sections was stained with the antibody against MAP2, a microtubule-associated structural protein highly expressed in the neuron soma, dendrite and axon. It has been shown that MAP2 labels the neuronal cell soma, dendrites and axons with hippocampal neuron culture or N2a cell line cell culture models [13, 14]. MAP2 is also a good marker for axons in pathological studies, which has been applied successfully in rat brain traumatic axonal injury model studies [15–17]. Although some cell culture experiments suggested that MAP2 is a proximal-axon-restricted marker in cultured dorsal root ganglion sensory neurons[18], however, if looking at the published data carefully, MAP2 expression is not confined to the proximal axons but also extended along the length of axons. MAP2 staining pattern might be very sensitive to experiment conditions given its profound changes during neurodegeneration. Different *in vivo* or *in vitro* experiment conditions such as different cell types, cell culture and cell growth conditions or staining of different MAP2 isoforms might explain the heterogeneity of the MAP2 staining results. In our immunostaining experiments on AD brain pathological sections and control brain sections, MAP2 stained the neuron soma and axons strongly and was clearly expressed in Layer VI axon bundles and in the axons in the white matter (Figure 1, 2 and Supplementary Figure 3), which proved that MAP2 is a useful marker for axon analysis in human brain pathology studies. The presented image showed that there was a significant decrease of signals of MAP2-stained axons (MAP2-positive axon numbers decreased more than half, at around 53.4% when examining from the longitudinal orientation (the top panel of Figure 2A). The counting from the transverse orientation image displayed even larger decrease around 79.8% of axon MAP2 signals (the bottom panel of Figure 2A). Double immunostaining with Aβ antibody showed that MAP2 expression inversely correlated with Aβ staining on the axons in AD brain tissues (Figure 2B). In AD brain white matter, many axons presented a decrease of MAP2 protein while bearing significant Aβ staining. MAP2 staining on Aβ-positive axons was often fragmented and diffusive. Within dense-core senile plaques, there is usually very little MAP2 immunostaining. The reduced MAP2 expression in amyloid-laden axons in the white matter could also be observed when using amyloid blue autofluorescence as an axon amyloidosis marker (Figure 2C). It is estimated, based on one set of experiments, Aβ immunohistochemistry labeled proximal axons around 44.2% by counting the average number of Aβ-labeled axons per axon bundle at transverse orientation (8.44±4.10, N=16) and divided by the average axon bundle axon number of 19.09 in control samples, which we obtained earlier. Axons with amyloid blue autofluorescence showed much weaker MAP2 intensity (0.36±0.07, N=100) comparing to axons without amyloid blue autofluorescence (N=25). In another word, amyloid-laden axons lost MAP2 expression at an average rate of 64% in the white matter. Besides the decrease of MAP2 protein in axons, reduced MAP2 immunostaining intensity in the neuron soma was also observed in neurons with intracellular Aβ (Supplementary Figure 5), suggesting that Aβ could have a deleterious effect on MAP2 expression both in the axons and in cell bodies.

**Figure 2.**
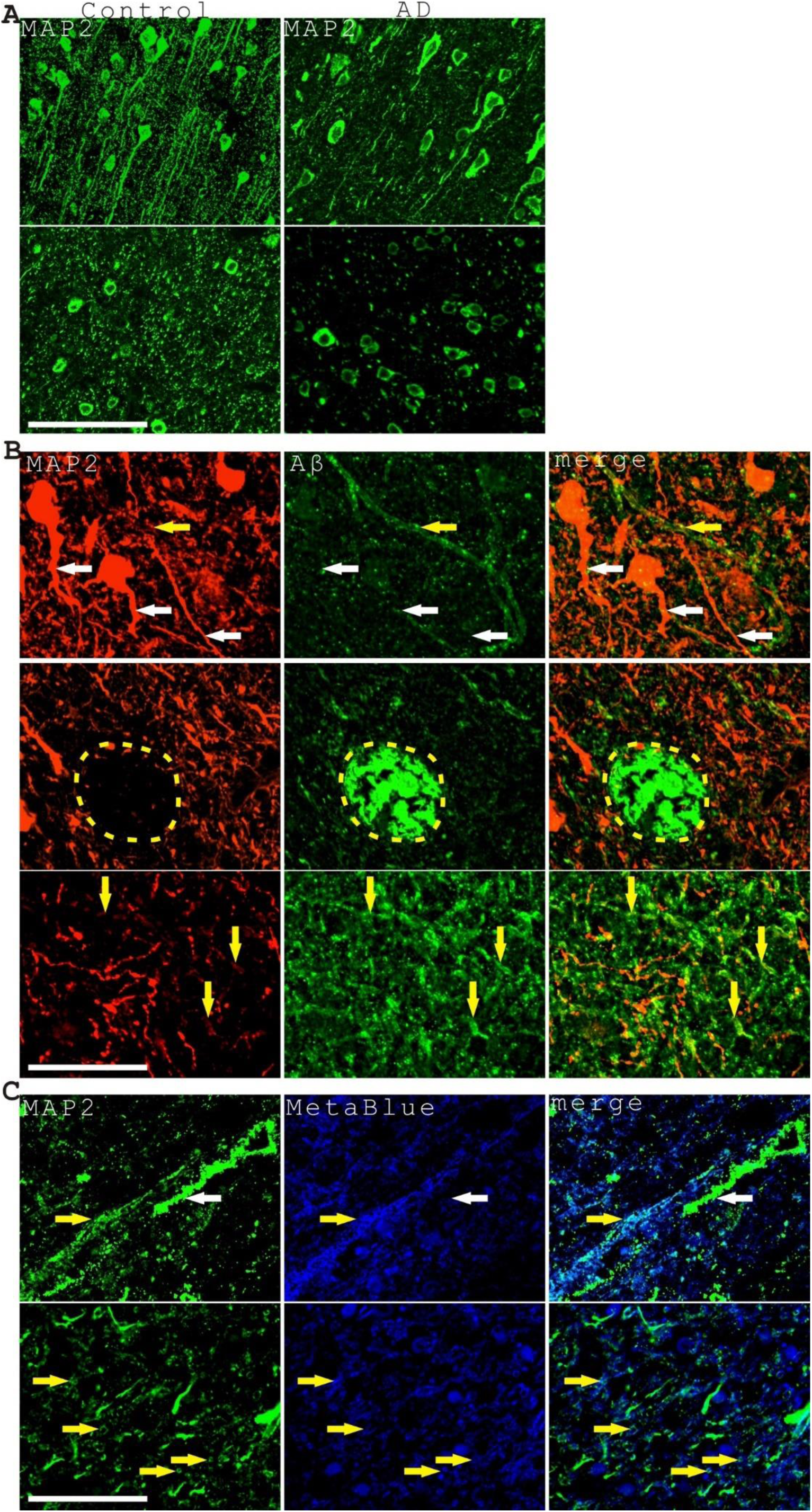
Axons in AD frontal brain tissues showed a decrease of axon structural protein MAP2. **A**. At longitudinal (top panel) and transverse orientation (bottom panel), a significant decrease of MAP2-positive axons in AD brain tissues comparing to control brain tissues was observed. The top panel showed that there were 26 long MAP2-positive axon fragments in the control sample while only 12 long MAP2-positive axon fragments in the AD sample, indicating a 53.8% decrease. The bottom panel showed there were around 391 strong MAP2-positive axon dots in the control brain while there were around 79 strong MAP2-positive axon dots in AD brain, which indicated a 79.8% decrease. Scale bar, 100 μm. **B.** Four representative images showed that the decrease of axonal MAP2 signals was correlated with increased Aβ immunostaining in axons in AD brain tissues. The top three images showed axons in the grey matter with greatly reduced MAP2 staining (amyloid-laden axons were indicated with yellow arrows while unaffected axons were indicated with white arrows). The yellow dashed-lines in the third image circled a dense-core senile plaque with barely detectable MAP2 immunostaining. The bottom image showed many Aβ-loaded axons with diminished MAP2 staining in the white matter. Scale bar, 50 μm. **C**. Two representative images showed greatly reduced axonal MAP2 staining in amyloid-laden axons labeled with amyloid blue autofluorescence in the white matter. The top panel showed axons mostly in the longitudinal direction and the bottom panel showed axons mostly in the transverse direction. The yellow arrows indicated the amyloid-loaded axons while the white arrow indicated a control axon without blue autofluorescence. Axons with amyloid blue autofluorescence showed much weaker MAP2 intensity (0.36±0.07, N=100) comparing to axons without amyloid blue autofluorescence (N=25). Scale bar, 50 μm.

### Lysosomal destabilization was detected in AD frontal brain tissues

We hypothesized that the decrease of MAP2 protein in axons might be due to some faulty protein degradation events although other mechanisms such as the reduction of MAP2 mRNA transcription or protein synthesis are also possible. Lysosomes and proteasome pathway are major pathways of protein degradation. Since we observed that Aβ-positive axons appear to contain many vesicles, we stained the sections with lysosomal enzyme marker Cathepsin D. Although previously we identified some weak extracellular Cathepsin D staining coming together with senile plaque Aβ aggregates, we also observed intracellular neuronal lysosome Cathepsin D staining clearly[3]. However, neuronal Cathepsin D staining was not homogeneous. Instead, we frequently observed abnormal Cathepsin D staining in the senile plaque regions (Figure 3). Normal lysosomal staining showed granule-like patterns with strong Cathepsin D signals. Conversely, nearly half of the lysosomal compartments in senile plaques showed abnormal Cathepsin D staining with enlarged, diffusive and weak staining patterns, indicating lysosomal destabilization, which has also been studied in previous literatures[19–21]. A weak but detectable Aβ staining could be observed in these abnormal lysosomal compartments. We measured the sizes and intensities of lysosomal compartments within senile plaque regions comparing to lysosomes outside of senile plaque regions. The lysosomal compartments (including all Cathepsin D-labeled compartments) within the senile plaques are roughly 4.32±1.99 μm (N=122) in size, which is 3.65±1.68 (Mann Whitney test, p<0.001) times of control lysosomes outside of plaque regions (1.18±0.39 μm, N=60) and characterized by considerably weaker Cathepsin D staining intensity (0.76±0.3) (Mann Whitney test, p<0.001). In AD axons, Aβ and lysosomal enzyme marker Cathepsin D are highly co-localized, showing the predominant presence of axonal Aβ in the lysosomes. We also detected diffusive and weaker Cathepsin D staining patterns along the length of Aβ-positive axons, suggesting axonal lysosome destabilization was happening (Figure 3D-F). Control cellular lysosomes far away from senile plaques had strong, clear granule-like Cathepsin D staining patterns (Figure 3G). Additional evidence showing axonal amyloidosis associating with unusually diffusive Cathepsin D staining was presented in Supplementary Figure 6. Amyloid blue autofluorescence was used as an amyloidosis marker in this figure instead. However, lysosome destabilization was not limited to axons, when looking at intracellular lysosomes more carefully, many neural cells also revealed heterogeneous Cathepsin D staining patterns, which were presented further in Figure 4. To summary, we think that the destabilized lysosomes might be related to the decrease of MAP2 protein in the axons.

**Figure 3.**
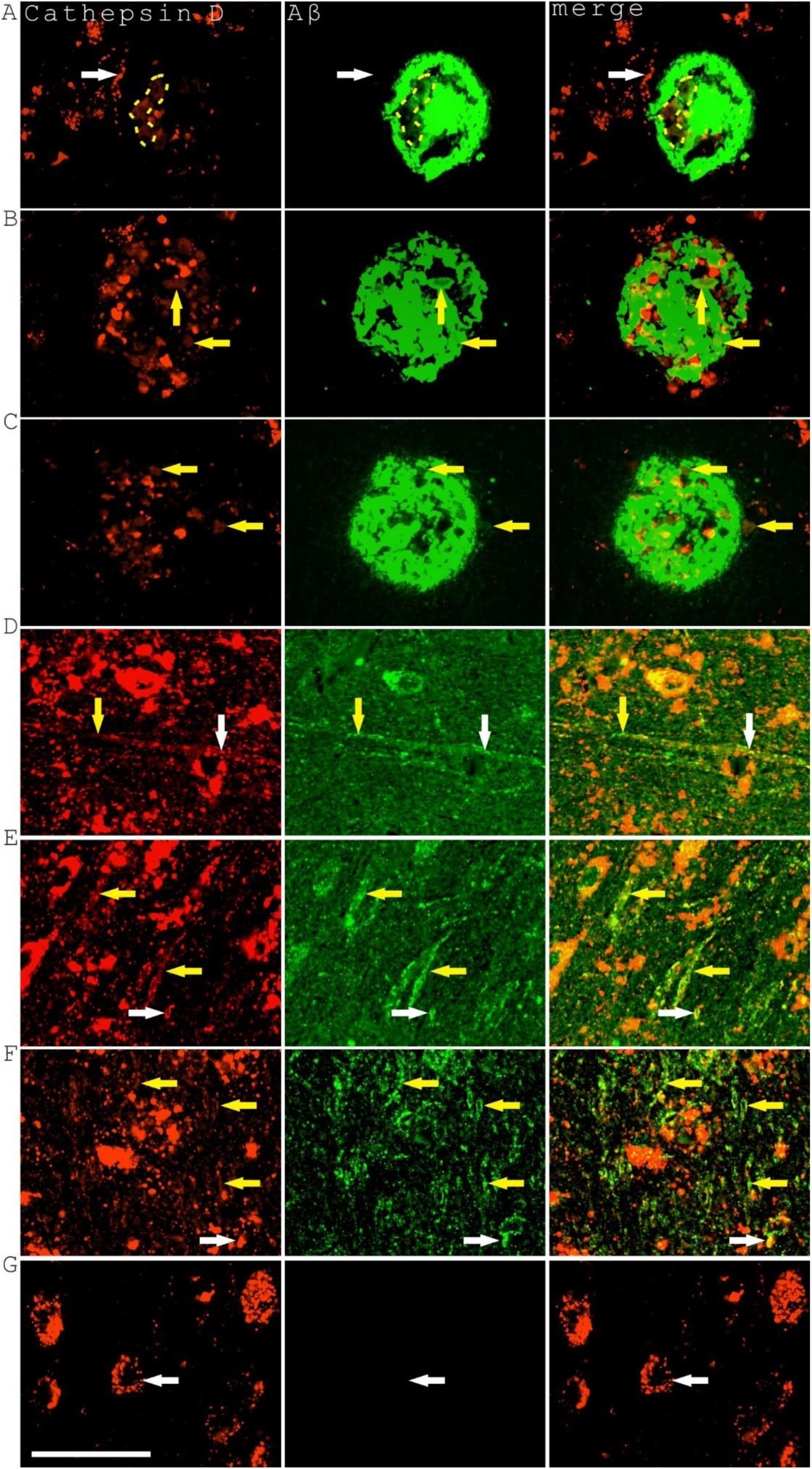
Lysosomal destabilization was common in AD frontal brain tissues. Three representative plaques were shown associating with lysosome destabilization (**A-C**). The top plaque was a typical dense-core plaque while the bottom two plaques were more diffusive. All three plaques associated with enlarged and diffusive Cathepsin D staining patterns (yellow dashed lines or yellow arrows), a character of lysosome destabilization and lysosome permeabilization comparing to typical, granule-like strong lysosome staining with focal Cathepsin D positivity (indicated with white arrows in the top panel). Weak Aβ staining could be observed in these abnormal lysosomal compartments. (**D-F**) showed three representative images illustrating the diffusive Cathepsin D staining in the Aβ-positive axons (indicated with yellow arrows). At the same time, the co-localization between Aβ and Cathepsin D in lysosomes was abundantly observed (indicated with white arrows). It should be noted that these three pictures were taken with longer exposure settings in order to show the weak axonal Aβ and Cathepsin D staining comparing to other pictures. **G**. Intracellular lysosomes away from amyloid plaques showed mostly clear, strong and granule-like lysosomal Cathepsin D staining (indicated with white arrows). Scale bar, 50 μm.

**Figure 4.**
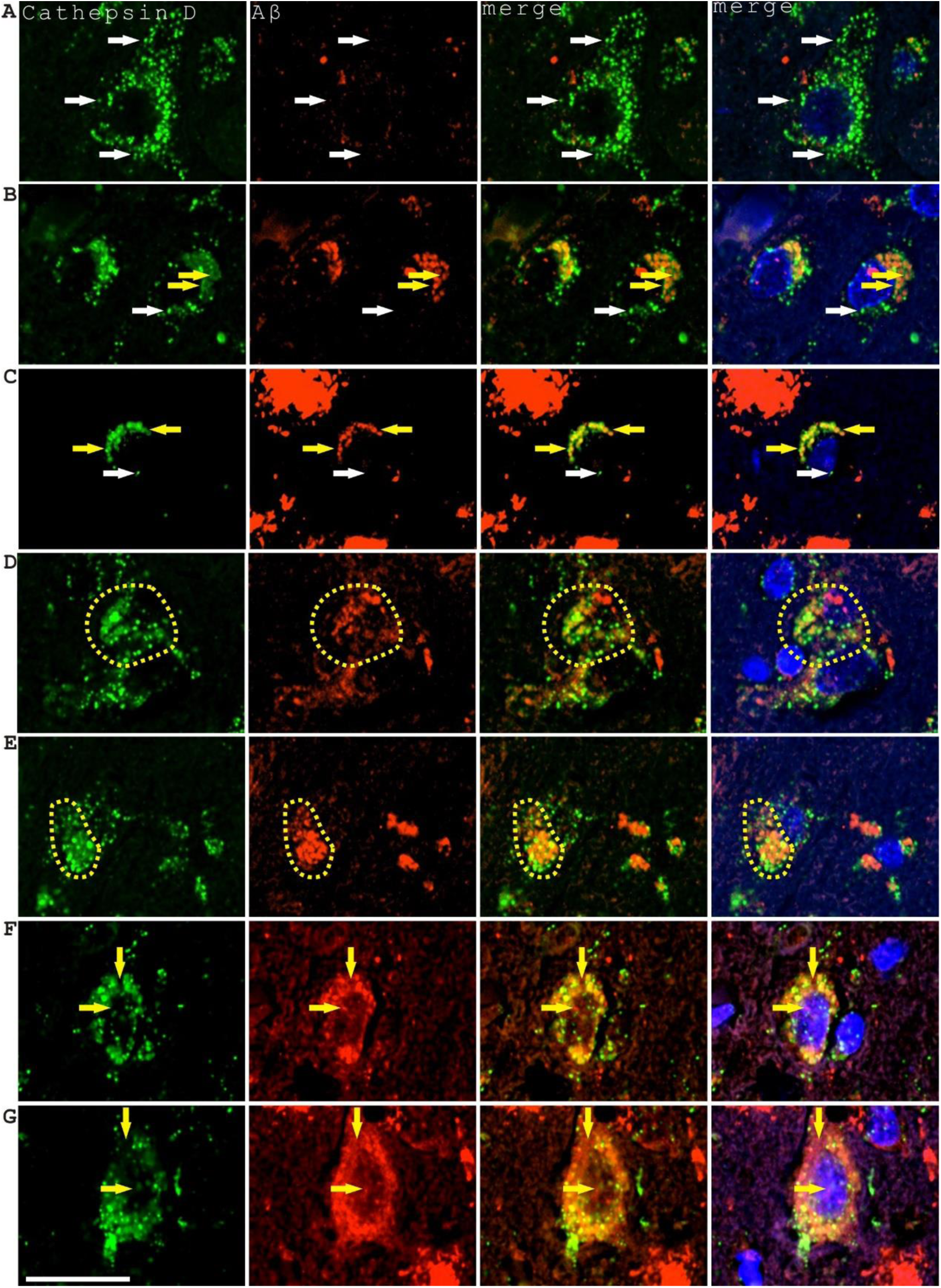
Lysosomal destabilization highly associated with Aβ in the cell bodies of neural cells in AD frontal brain tissues. **A.** A control neural cell with little Aβ staining showed mostly normal lysosome Cathepsin D staining patterns (indicated with white arrows). **B, C.** Intracellular Aβ associated with lysosome destabilization, which could be identified at single lysosome levels with significant lysosome enlargement and diffusive staining (indicated with yellow arrows) while control lysosomes indicated with white arrows showing compact and bright Cathepsin D staining. **D, E** showed two representative images indicating that intracellular Aβ expression associates with lysosome enlargement, clustering and diffusive staining. **F**, **G** showed two representative images of neural cells containing not only lysosomal Aβ and Cathepsin D but also extra-lysosomal Aβ, nuclear Aβ and Cathepsin D. The vertical arrows indicated Aβ staining outside of Cathepsin D-stained areas while the horizontal arrows indicated nuclear Aβ and Cathepsin D staining. Quantitative measurement showed that Aβ-containing cell body lysosomes (N=30) had an average size of 1.63±0.27 μm, which is 2.23±0.38 times of control lysosomes without Aβ staining (0.73±0.09 μm, N=12) and an average Cathepsin D intensity of 0.32±0.12 time of control lysosomes. Scale bar, 25 μm.

Lysosomes are abundantly present in the neuronal soma. If lysosome destabilization is happening surrounding senile plaques, we should also expect to see some destabilized lysosomes in the cell bodies. Not surprisingly, we did observe frequent lysosomal destabilization in Aβ-containing neural cells (Figure 4 and Supplementary Figure 7). We use neural cells with little Aβ staining as internal controls for unaffected lysosomes (Figure 4A). The control lysosomes appeared normal-looking with punctuate, compact staining of Cathepsin D. In cells with abundant Aβ signals, lysosomes became enlarged, clustered and their Cathepsin D staining became diffusive. The enlargement of lysosomes filled with Aβ could be observed at single lysosome level (yellow arrows in Figure 4B, 4C). The Aβ-loaded and enlarged lysosomes often clustered with each other to form large lysosomal domains with diffusive Cathepsin staining (Figure 4D, E). Quantitatively, Aβ-containing lysosomes (N=30) had an average size 1.63±0.27 μm, which is 2.23±0.38 times (the difference is statistically significant with p<0.001, T-test) of control lysosomes with no Aβ staining with the average size of (0.73±0.09 μm, N=12) and an average intensity of 0.32±0.12 times (p<0.001, Mann-Whitney test) of control lysosomes, demonstrating that Aβ-associated lysosome destabilization is a distinctive phenotype of intracellular amyloidosis. We also observed that Aβ staining sometimes showed up at extra-lysosomal locations in the cytoplasm or even in the nucleus. Occasionally, we also observed Cathepsin D staining in the nucleus (Figure 4F, G). The existence of Aβ and Cathepsin D in locations other than lysosomes might also be related to lysosome destabilization and leakage, which theoretically could relocate lysosomal contents to other cellular compartments.

The above result strengthened the concept of lysosome destabilization in AD as also reported from previous studies [20–23], which can be regarded as a consensus in the field. However, whether abnormal lysosomes produced Aβ or Aβ induced lysosomal destabilization is still a debated issue. Nevertheless, we could tackle this difficult question in some way by studying the molecules interacting with Aβ. Previously, we established a linkage between Aβ senile plaque formation and intravascular hemolysis, vascular degeneration and microaneurysm rupture[3, 9]. The relationship between microhemorrhage and Aβ senile plaque information has also been explored in other published investigations[24–30]. Thus, Aβ might interact with hemolysis or vascular markers either directly or indirectly, which should be reflected with the co-distribution of Aβ with hemolytic or vascular markers. We analyzed the intracellular co-expression of a hemolysis-related marker HBA (alpha Hemoglobin) and two vascular-related markers ACTA2 and ColIV with Aβ (Figure 5). All three markers showed good co-localization with intracellular Aβ in lysosome-like structures. HBA and ACTA2 signals as well as Aβ signals were also frequently observed in the nucleus (Figure 5A). When we performed HBA and Cathepsin D double-immunostaining on AD sections, the co-distribution of HBA signals with destabilized lysosomes as indicated with enlarged and clustered lysosome compartments and diffusive Cathepsin D staining was also observed (Figure 5B). This indicated that the protein partners interacting with Aβ such as HBA also associated with lysosome destabilization in neural cells.

**Figure 5.**
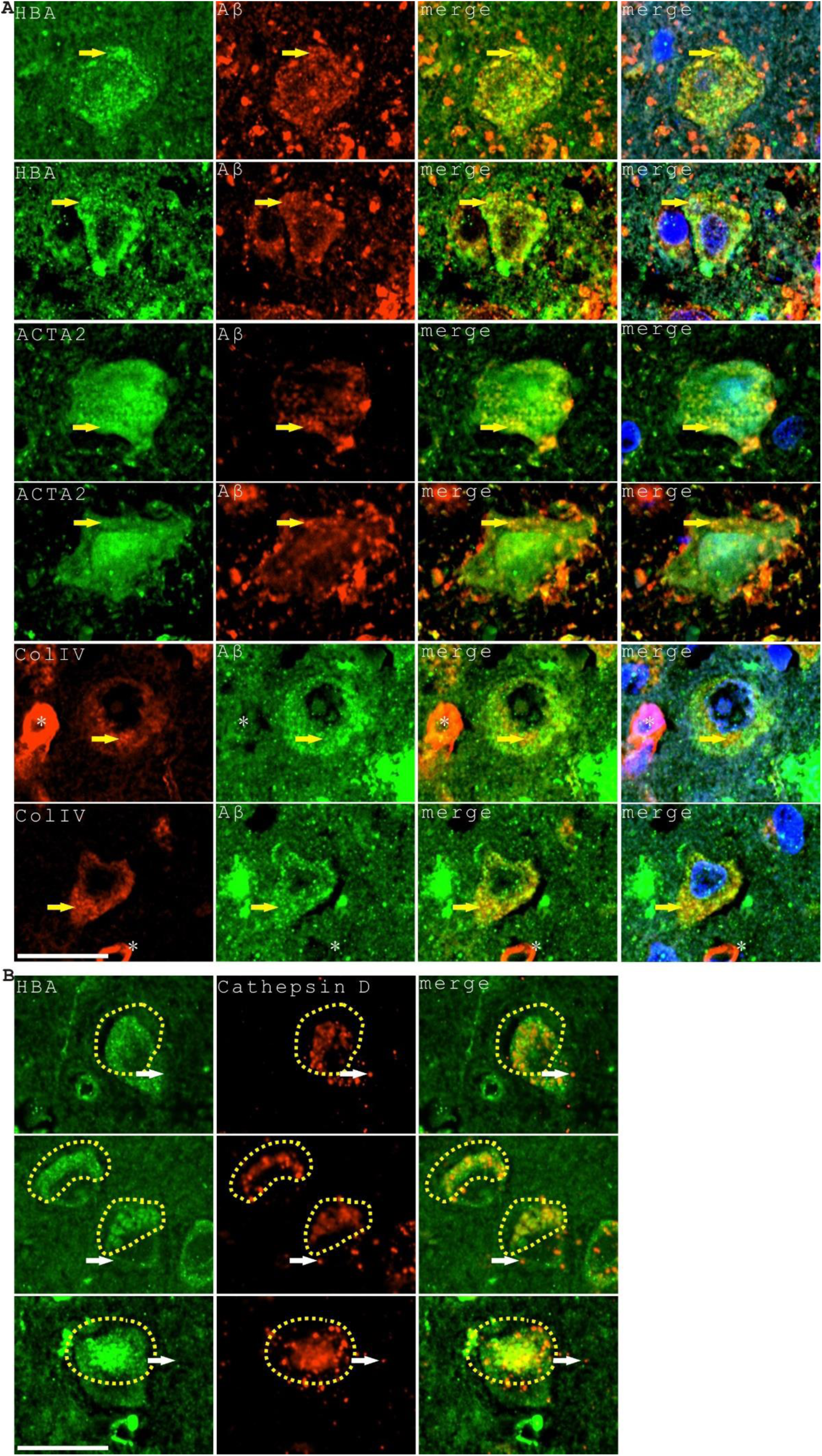
Blood and vascular proteins co-existed with intracellular Aβ in neural cells in AD frontal brain tissues and might also be related to lysosome instability. **A**. Representative images showing the co-expression of blood related marker HBA, vascular related markers ACTA2, ColIV and Aβ. The co-localization of markers was indicated with yellow arrows. Two small blood vessels were indicated with asterisks in the ColIV staining panel. **B**. The intracellular HBA expression domains overlapped with the region of lysosomal destabilization indicated with enlarged and clustered lysosomal compartments and diffusive Cathepsin D staining (marked with yellow dashed circles). Control HBA-unaffected lysosomes were indicated with white arrows. Scale bars, 25 μm.

### Destabilized lysosomes existed in dystrophic neurites

It is clear that there is a lysosome destabilization phenotype in neuronal cells associating with intracellular Aβ from studies above. Since we focused more on the axonal phenotype in AD, to further verify if destabilized lysosomes were linked to axon degeneration, we stained the tissue sections with phosphorylated Tau (phos-Tau) antibody, a well-known axon dystrophy marker in Alzheimer’s disease, and the lysosomal membrane marker Lamp2. The localization of abnormally diffusive lysosomal Lamp2 staining in phos-Tau-positive dystrophic neurites, neurofibrillary tangles, axons and neuropil threads was observed (Figure 6A). In addition, the abnormally diffusive lysosomal Lamp2 staining in phos-Tau-positive dystrophic axons was also abundantly detected in the white matter (Figure 6A, bottom panel). Neuropil thread is a collective term traditionally used for phos-Tau staining in both axons and dendrites. We noticed that, across broad brain regions, neuropil thread phos-Tau staining was more abundant than phos-Tau staining of the tangles and senile plaque dystrophic neurites (Supplementary Figure 8). phos-Tau positive neurites in AD tissues could be classified with primary dystrophic neurites at an average diameter of 1.56±0.37 μm (N=40) or secondary dystrophic neurites with an average diameter of 0.40±0.09 μm (N=30) based on their diameters, which might reflect the different size of axons or dendrites. When we stained AD brain sections with both phos-Tau and Aβ antibodies, the co-localization between phos-Tau and Aβ was also detected in dystrophic neurites in senile plaques, neurofibrillary tangles, neuropil threads and axons (Figure 6B). However, Aβ staining in dystrophic neurites, tangles and axons was lower comparing to the strong Aβ staining in the senile plaques. These data confirmed that axon dystrophy connected with Aβ, Tau phosphorylation and lysosome destabilization.

**Figure 6.**
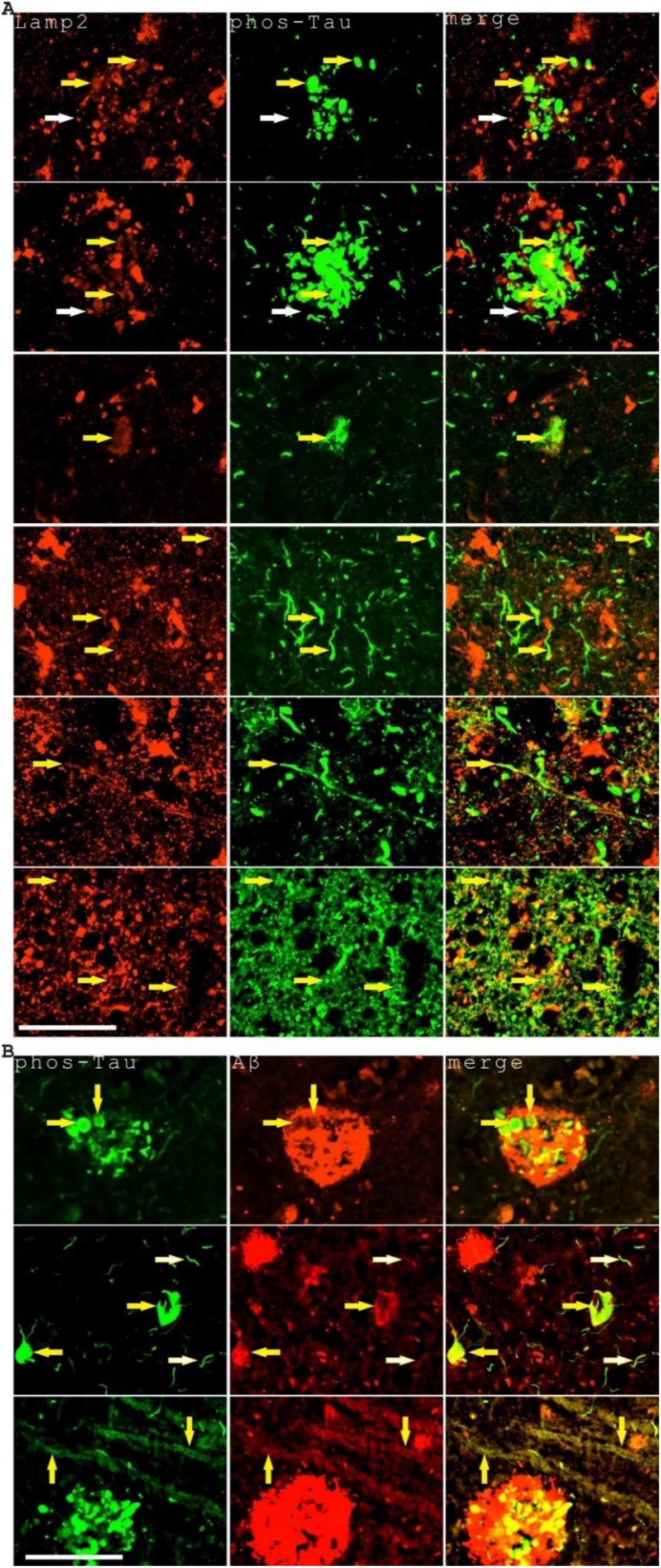
Destabilized lysosomes were located in AD frontal brain dystrophic neurites, tangles, axons and neuropil threads, additionally there is a co-distribution between phos-Tau and Aβ in dystrophic neurites, tangles, neuropil threads and axons. **A**. Abnormal lysosome Lamp2 staining (indicated with yellow arrows) was detected in peri-plaque dystrophic neurites (top two panels), a representative tangle (the third panel), a representative longitudinal axon (the fourth panel) and white matter axons mostly in the transverse direction (the bottom panel) marked with phos-Tau immunostaining. We observed long, diffusive but relatively weaker Lamp2 staining along the length of axons and neuropil threads, highlighting the intricacy of this defect. The bottom three picture was taken with longer exposure time settings in order to show the relatively weak signals in neuropil threads and axons. Control lysosomes not affected by Tau phosphorylation were indicated with white arrows. Scale bar, 50 μm. **B**. phos-Tau-positive dystrophic neurites (yellow arrows, top panel), tangles (yellow arrows) and neuropil threads (white arrows) (middle panel) and axons (yellow arrows, bottom panel) were also Aβ-positive. Scale bar, 50 μm.

### Axon degeneration in AD bears the markers of hemorrhagic insults

It is not very clear why cells and axons in AD tissues are degenerating. A lysosome destabilization mechanism was proposed initially based on cell culture studies [20, 21]. However, in the *in vivo* situations, Aβ alone might not be fully responsible for the degenerating effect since Aβ is interacting with many other proteins. As our previous research suggested that senile plaques are formed by blood Aβ leakage into the brain upon microaneurysm rupture, we hypothesized that axon damages in AD tissues might be also associated with many blood-related proteins. We checked the axon pathology with a variety of blood or plasma-related markers. The results showed that enlarged axons in AD brains indeed carried blood and plasma markers such as ApoE, HBA, HbA1C and Hemin (Figure 7A-D). Overt acute hemorrhage was rarely observed surrounding these affected axons, suggesting that the intake of hemorrhagic markers in these axons is likely a chronic event. It appears that the presence of blood markers in the axons is not depending on the spatial closeness to the senile plaques. In addition, the distribution of hemorrhagic markers is not always quantitatively proportional to the intensity of Aβ in AD brain tissues. Sometimes there is even stronger staining of hemorrhagic markers in the axons than in the surrounding senile plaques, which is likely because of the differential enrichment or metabolism of amyloid protein complexes in axons vs. senile plaques (an example was shown in the top panel of Figure 7D with more Hemin staining in the axons than in the adjacent senile plaque). Moreover, axon amyloidosis is associated with the expression of Sortilin1, another endosomal/lysosomal marker (Figure 7E), which has also been implicated as a major receptor for ApoE and a component of senile plaque[31, 32]. Since intracellular Aβ mostly existed in the lysosomes as shown in Figure 3, the above data also suggested that intracellular ApoE, HBA, HbA1C and Hemin primarily located in the lysosomes. An overview image of widespread HBA and Aβ expression in the white matter axons was further presented in the Supplementary Figure 9. Additionally, when we stained the sections with red blood cell related histological stains such as Alizarin red and Rhodanine[3], we could also see many axons in AD brain tissues positively stained by these stains (Supplementary Figure 10). Actually, AD white matter is broadly stained with Alizarin red and Rhodanine, similarly as HBA, showing an extensive effect of hemorrhagic markers on the white matter axons. All these analyses indicated that the axons not only were enriched for Aβ but also enriched for many hemorrhage-related markers.

**Figure 7.**
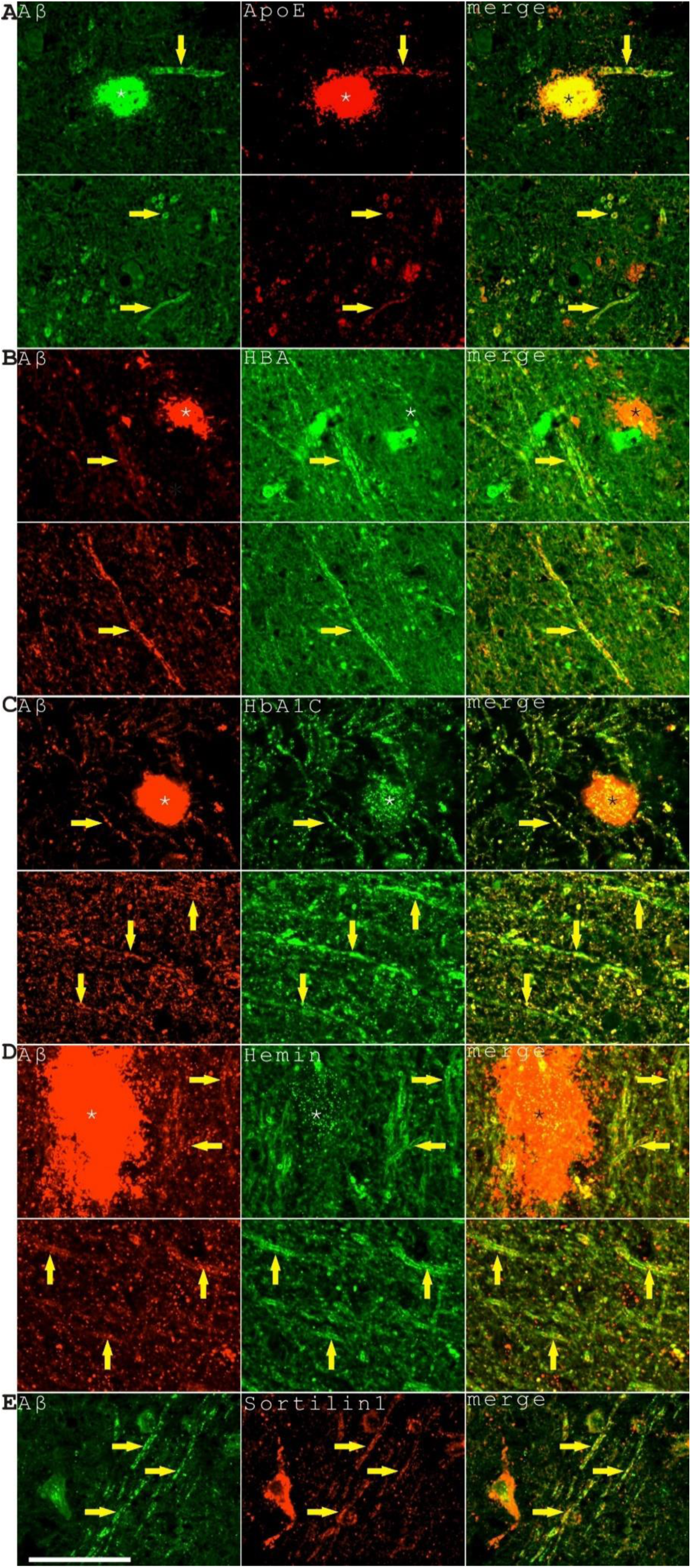
Axon degeneration in AD bears the markers of hemorrhagic insults in AD frontal brain tissues. Four blood-related markers (ApoE, HBA, HbA1C, Hemin) all showed positivity in enlarged axons labeled with Aβ immunostaining, that can be adjacent to senile plaques (top image in each panel) or not associated with senile plaques (bottom image in each panel) (**A** to **D**). Degenerating axons were also positive for an endosomal/lysosomal marker Sortilin1 (**E**). The affected axons were indicated by arrows while the senile plaques were indicated with asterisks. Scale bar, 50 μm.

### Under rare occasions, axon breakages were observed in AD brain tissues

Whether the axonal degeneration defects will ultimately lead to axon breakages in Alzheimer’s disease is an important question to be addressed. We checked hundreds of images containing Aβ-stained axons. We initially focused on axons traveling in the longitudinal orientation inside of senile plaques since the axons in the transverse orientation were difficult to be individually identified. We observed that many axons passing through the senile plaques were Aβ-positive and enlarged but they did not break (Supplementary Figure 11). However, we did find a few examples indicating that the axons could truly be broken at locations outside of senile plaques. Using Aβ and AGE antibody as the markers for amyloidosis-affected axons, broken axons with gaps up to 38.4 μm in width could be detected (Figure 8A). What caused these rare axon breakages is not immediately clear. On the other hand, we observed that long stretches of amyloid-loaded vesicles with large spheroid formation on the axons caused dramatic dystrophy to a degree that the axon morphology could no longer be recognized (Figure 8B). The long stretch clustering of vesicles with spheroid formation was likely due to lysosome clustering as indicated by the endosome/lysosomal marker Sortilin1 (Figure 8C). It is possible that the continuous enlargement of destabilized lysosomal compartments across long distance might finally result in axon breakages in some cases.

**Figure 8.**
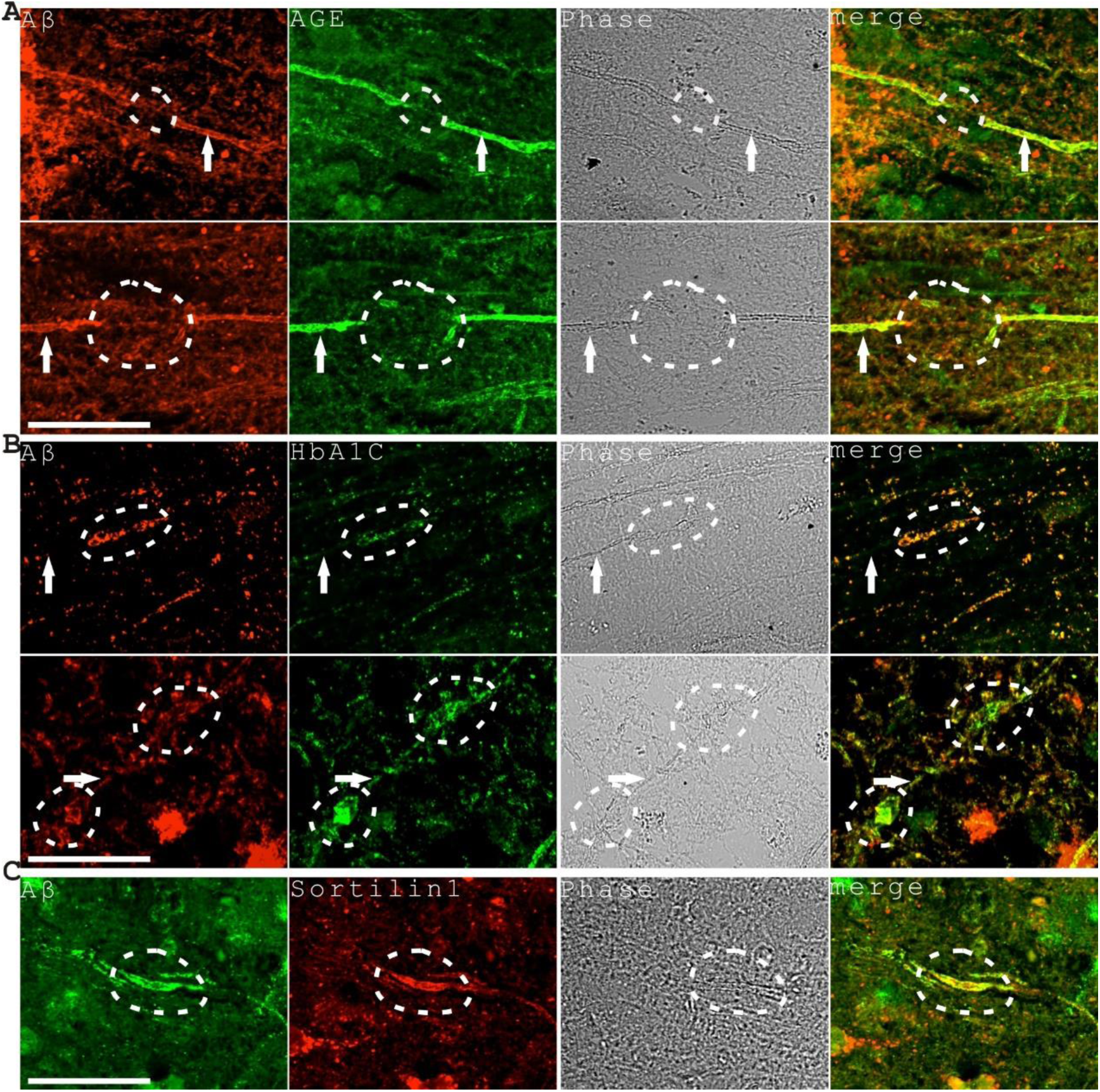
Under rare occasions, axon breakages were observed in AD frontal brain tissues. **A**. Two broken axons were identified with AGE and Aβ immunohistochemistry in AD brain tissues. The axonal gap in the top image was measured around 24.4 μm while in the bottom image was measured around 38.4 μm. The broken regions were marked with dashed lines while the parental axons were indicated with arrows. **B**. Two examples of damaged axons associated with large vesicle clusters and spheroid formation enriched for amyloid markers Aβ and HbA1C. The long stretch of clustered vesicles (indicated with dashed lines) in the top panel measured around 32.5 μm in length. The maximal width of the two large spheroids (indicated with dashed lines) in the bottom panel reached 6.2 μm (top-right position) and 8.3 μm (bottom-left position) respectively. The affected axons were indicated with arrows. **C**. A long axonal region (indicated with dashed lines) with large spheroid formation associated with endosomal/lysosomal domains marked with Sortilin1 immunostaining. This region was measured around 26.1 μm in length. The maximal width of the spheroid measured around 4.1 μm. Scale bars, 50 μm.

## Discussion

The reason that axonal Aβ staining was not clearly observed in our previous studies[3, 9] is likely because of the relatively small size of axons and the larger and stronger senile plaque Aβ staining camouflaging the fainter staining of axonal Aβ. The widespread axonal enlargement associated with Aβ deposition might have a significant influence on AD pathology. It has been shown that in the natural animal species, axons with larger diameter have higher transduction speed, for example, in the case of the squid giant axon[33]. However, in the case of AD, the enlarged axons were not natural but instead pathological, which were filled with toxic Aβ, hemorrhagic proteins and destabilized lysosomes. In addition, lysosomal leakage can induce axonal structural damages and increase axolemmal permeability, which might further damage the myelin sheath. Both human and mouse studies indicated that axon diameter increases with aging and cognition decline[34, 35]. There is also a reduced velocity of action potential transmission in AD patients and AD-model transgenic mice[36, 37]. Diffusion tensor imaging studies also indicated that structural damage and disconnection in the white matter is prominent in AD [38, 39]. Axonal amyloidosis and enlargement in AD are likely linked to transduction velocity decrease and white matter damage phenotypes. Besides the basic axonal enlargement phenotype in AD, spheroid formation is also frequently observed. Previous studies showed that spheroid formation on axons also slows down the speed of action potential transduction[40, 41]. All these studies presented strong evidence that there are serious axonal defects in AD with transduction speed decline as a central phenotype. How and to what extent Aβ deposition in enlarged axons impacts on the whole brain connectome requires further large-scale quantitative analysis and *in vivo* animal and human studies.

Axonal amyloidosis additionally induces structural changes such as a decrease of MAP2 protein, which might be related to Aβ toxicity in inducing lysosomal destabilization. It has been shown *in vitro* that extracellular Aβ was taken up and concentrated in lysosomes and induced lysosome destabilization and permeabilization[20, 21]. Lysosome permeabilization has been studied in mouse AD models and was considered as the possible mechanism for neuronal cell death and senile plaque formation[23, 42, 43]. A previous study also identified abnormal protease-deficient lysosomes in the swollen axons surrounding senile plaques[22]. The study defined the lysosome defect as a lysosome maturation deficiency. In this study, we provided evidence that Aβ is linked to lysosome destabilization at single lysosome level and single axon level. We think that the axonal or cellular lysosomes were not intrinsically protease-deficient but rather be destabilized by Aβ and Aβ-associated hemorrhagic or vascular factors such as ApoE, Hemin, HbA1C, HBA, ACTA2 and ColIV. It is entirely possible that Aβ-associated factors are also affecting lysosome stability, maturation or lysosome pH besides Aβ itself. This study further supported our previous studies that the pathological Aβ peptide in AD tissues comes from an exogenous source, the hemorrhagic Aβ leakage, instead of coming from neurons themselves[3, 9]. Although it is possible that lysosome leakage might leads to neuronal cell death, we think lysosome leakage in the cells could illicit a complicate "lysosome leakage response" to counter-react the damaging effect of lysosome leakage. Our preliminary data did not confirm a straightforward increase of neuronal apoptosis with TUNEL assay as an apoptotic marker and MAP2 reduction as a marker for lysosome-leakage associated proteomic damage. It warrants more extensive investigations on the mechanism of "lysosome leakage response" in the future.

In our opinion, Aβ could be classified as a type of lysosomal “toxin”, which exerts its unique toxicity upon lysosomal intake and processing. The current study emphasized that lysosomal destabilization might not be due to Aβ alone since neuronal cell amyloidosis associated with the enrichment of multiple hemorrhagic or vascular markers such as ACTA2, ColIV, HBA, ApoE, HbA1C and Hemin as shown in Figure 5 and Figure 7, which are either major Aβ binding partners or major components of microhemorrhage [3, 9, 28, 44]. It is well-known that Hemin is a neurotoxic molecule by itself[45, 46]. Axonal amyloidosis also associated with Sortilin1 expression. Sortilin1 is an endosome/lysosome marker that has been identified as a major receptor of ApoE[31]. Axons might take up exogenous Aβ complexes through receptor-related mechanisms, independent of or in conjunction with the neural soma, by the relay chains starting from Aβ to ApoE then to Sortilin1. The intake of a large number of hemorrhagic proteins along with Aβ could possibly induce severe axonal proteostatic stress besides the toxicity effects of Aβ especially when the axon-soma connection might be compromised.

Both Aβ deposition and Tau phosphorylation are important components of AD pathology, however, how these two events connect is not clear. Previously, it has been suggested that Aβ and Tau co-localize in AD “synaptosomes”[47]. In this study, we provide another possible answer that Aβ and Tau are both connected to destabilized lysosomes. There is ample evidence that Aβ endocytosis induces lysosome destabilization[19, 20, 48]. There is also strong evidence that Aβ likely induces Tau phosphorylation[49, 50]. We show in this study that Tau phosphorylation associates with destabilized lysosomes in tangles, dystrophic neurites and also neuropil threads. It is not known whether Tau phosphorylation or lysosome destabilization is the upstream event. However, if taking into account for the MAP2 reduction in axons and the evidence in the literature that MAP2 competes with Tau for microtubule binding[51] and could prevent Tau aggregation[52], lysosomal destabilization might work upstream of Tau phosphorylation by chronically degrading its competitive inhibitor MAP2. The biological significance of lysosome destabilization in the neural soma in AD tissues is worthy of future investigations.

Under rare occasions, the breakage of axons has been observed in a small number of axons in our experiments. When axon breakages happen and are not repaired, Wallerian degeneration of axons might occur, which could result in axon loss. Since axon breakages with small gaps are difficult to detect, the actual axonal breakage events in AD brains might be more abundant than what we observed. How frequently the axonal breakages and Wallerian degeneration happen in AD brains requires further investigation. Previous *in vitro* study has shown that Aβ peptide actually induces axon degeneration precedes cell death in hippocampal neuron cell culture models[53], indicating a possible "dying back" mechanism of neuronal degeneration in AD. Both Wallerian degeneration and axon regeneration are affected by various genetic mutations[54–56]. To combat Alzheimer’s disease efficiently, firstly, we need to find ways to prevent microvascular blood leakage and axonal accumulation of Aβ and hemorrhagic proteins. Secondly, treatments that can specifically slow down Wallerian degeneration and facilitate axon repair, reconnection and regeneration are also likely strongly beneficial for AD patients.

## Acknowledgements

We want to thank the help from Dr. Ma Chao and Dr. Qiu Wenying for providing AD tissue sections from National Human Brain Bank for Development and Function, Chinese Academy of Medical Sciences and Peking Union Medical College, Beijing, China. Additionally, we want to thank Dr. Ming Yang from Shanghai Jiao Tong University, Dr. Xiangli Zhang from Shandong University, Dr. Peng Du and Dr. Lei Xia from Xinhua Hospital for valuable scientific discussions. Moreover, we want to thank Housheng Wang, Pingxin Liu, Xiaoyi Bao, and Jun Wang for excellent laboratory assistance.

## Supplementary Information

**Supplementary Figure 1.**
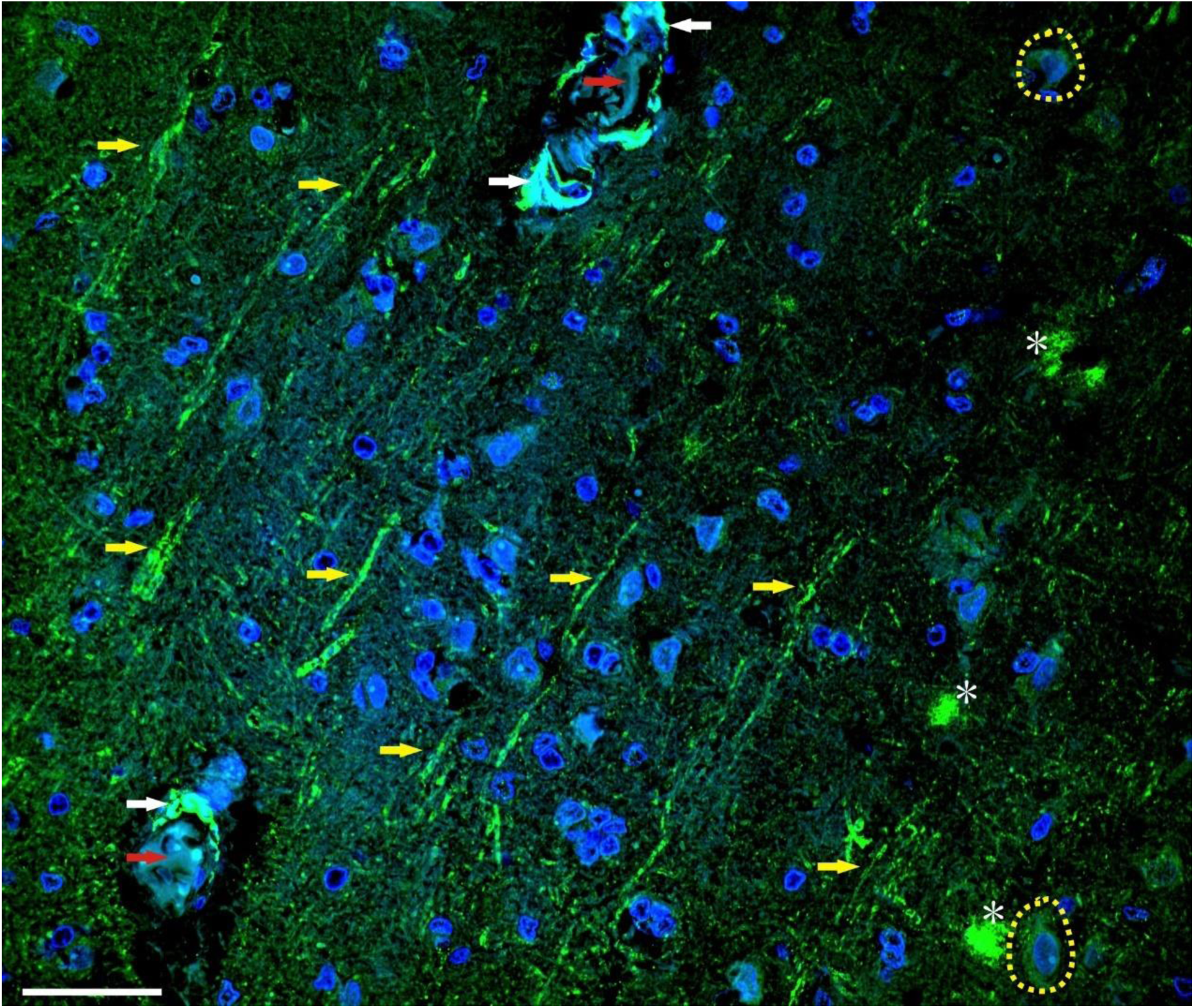
An overview picture indicated that Aβ expressed in axon bundles, senile plaques, CAA blood vessel walls, the blood vessel lumen and also in neurons in AD frontal brain tissues. Aβ was stained with green fluorescence and the nuclei labeled blue by DAPI. The yellow arrows indicated axon bundles, while white arrows indicated CAA blood vessels and the red arrows indicated the blood vessel lumens. Many indicated axons showed quite even enlargement of the axons. A few small senile plaques were indicated with asterisks and Aβ expression in neurons was indicated with yellow dashed circles. Scale bar, 50 μm.

**Supplementary Figure 2.**
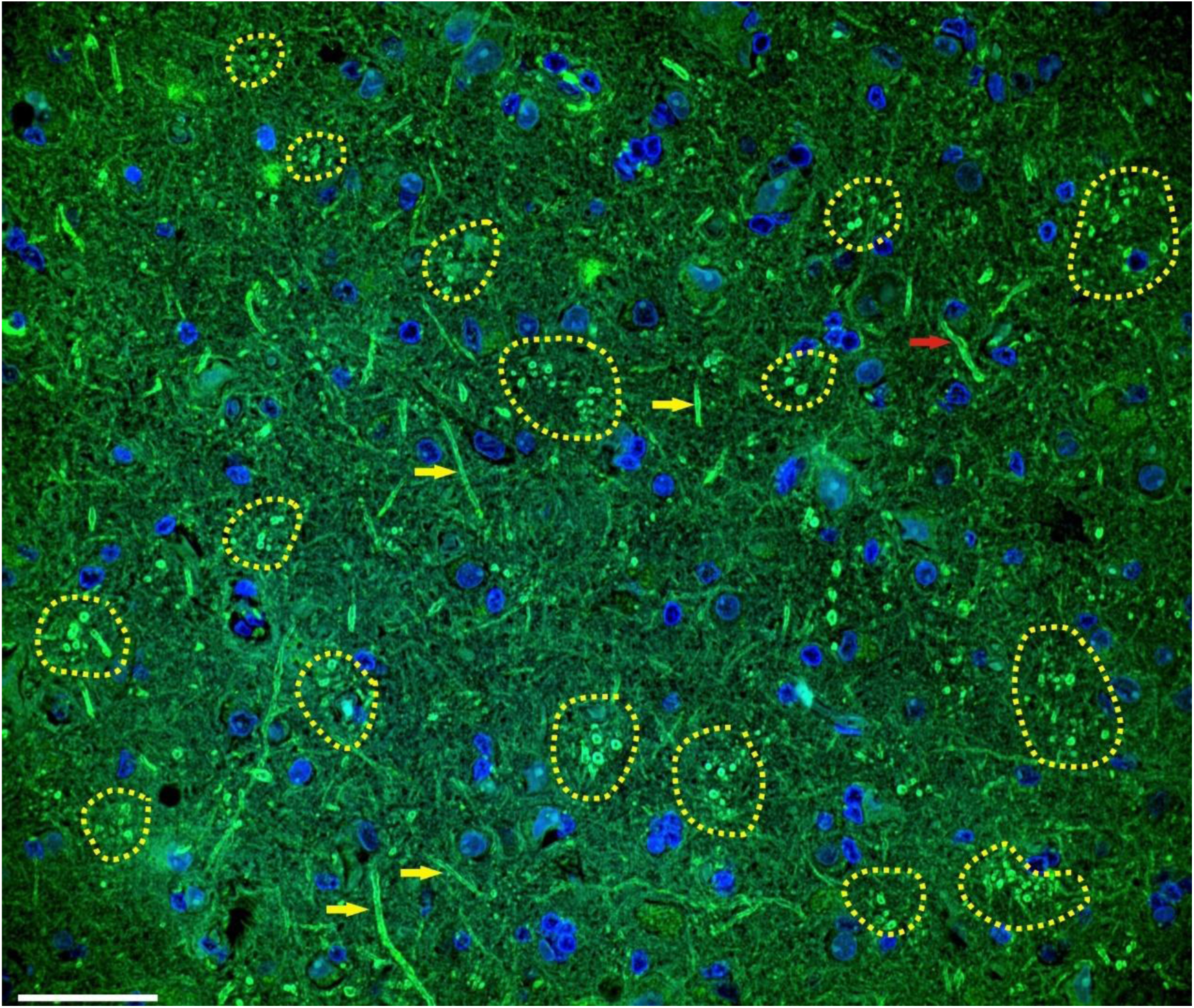
An overview picture illustrated many axon bundles with Aβ-positive axons in a transverse orientation in AD frontal brain tissues. Aβ was stained with green fluorescence and the nuclei labeled blue by DAPI. The yellow dashed circles indicated the axon bundles. Many axons at the longitudinal direction showed quite even enlargement of the axons as indicated with yellow arrows. An axon with spheroid formation was indicated with a red arrow. There are around 8.44±4.10 Aβ-positive axons on average in each axon bundle (N=16). Scale bar, 50 μm.

**Supplementary Figure 3.**
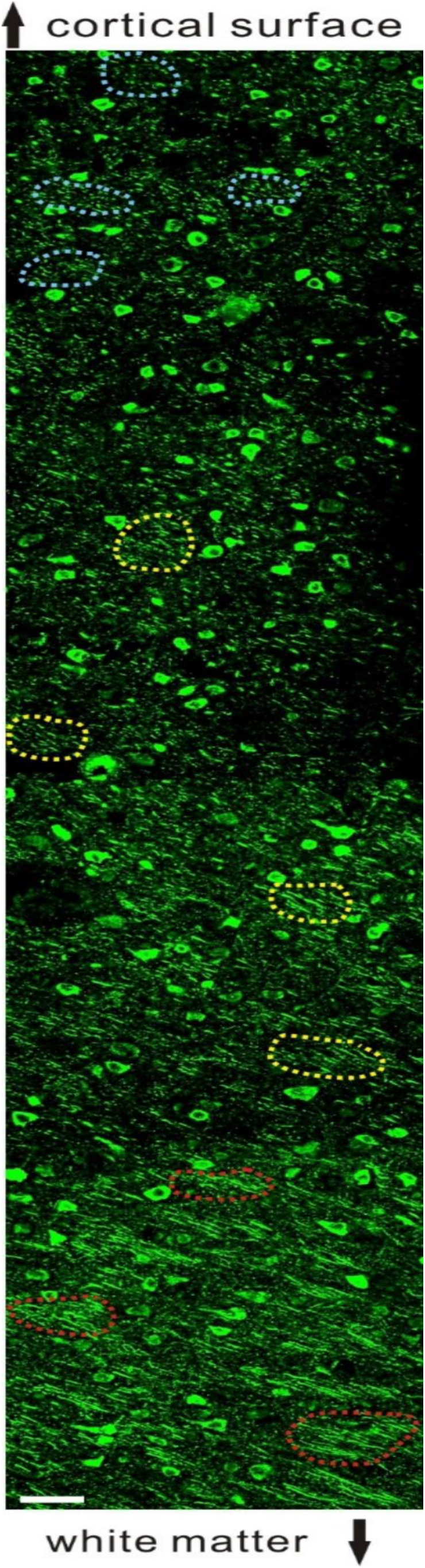
MAP2 antibody labeled intracortical axon bundles clearly in control frontal brain tissues. The orientation of this tissue slide is from the cortical surface (top) to the white matter (bottom). Axon bundles at both transverse (top, indicated with blue circles) and longitudinal orientation (bottom, indicated with red circles) could be observed in the same pathological section at different depth in the brain tissues. Axon bundles with intermediate length axon fragments could be observed in between (indicated with yellow circles). Not all but only representative axon bundles were indicated. These axon bundles has an average diameter of 61.61±9.88 um (N=11). Each bundle contains an average of 19.09±4.61 axons (N=11). Scale bar, 50 μm.

**Supplementary Figure 4.**
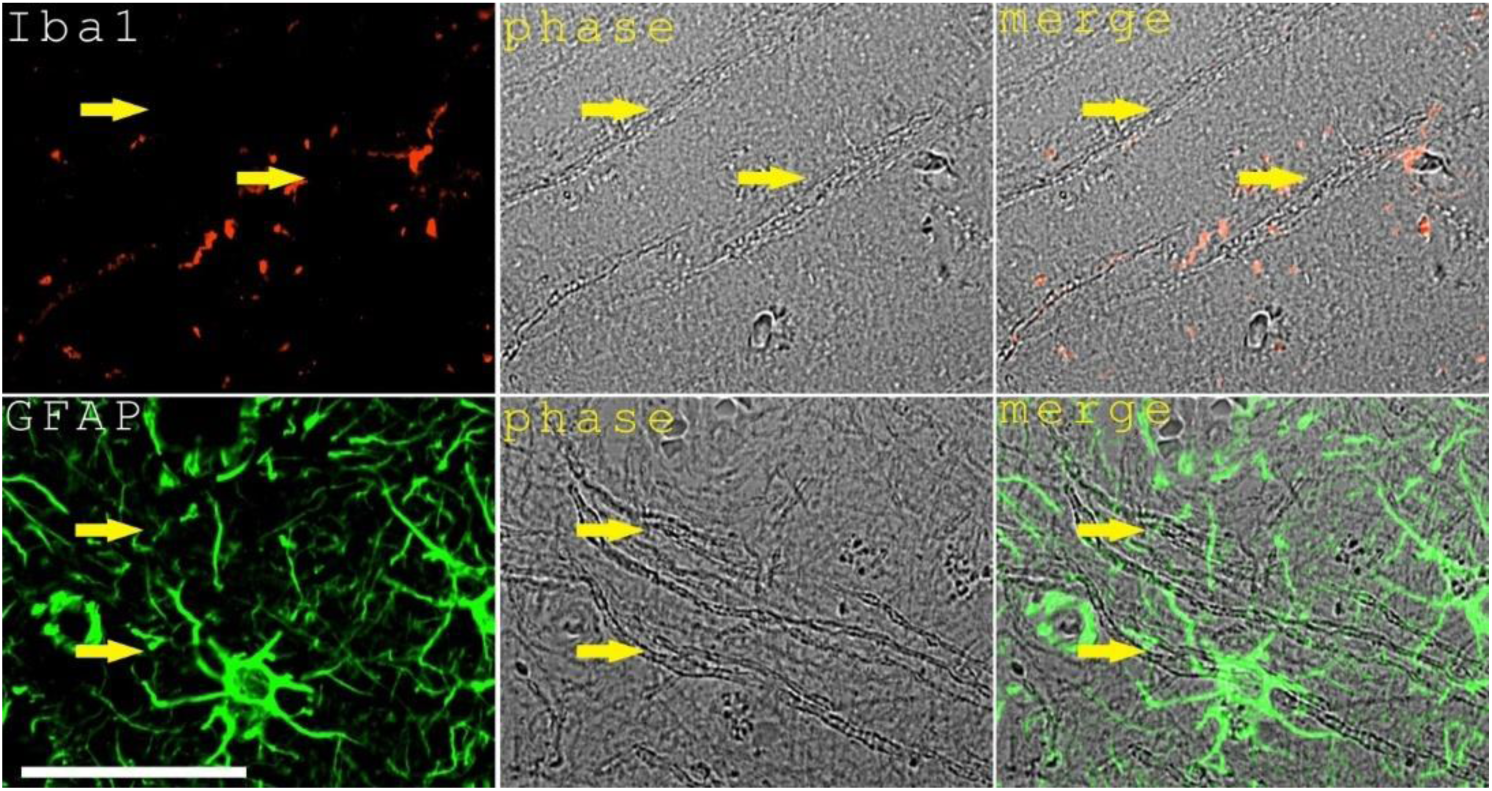
Enlarged axons in AD frontal brain tissues were not stained with microglial marker Iba1 (top panel with red fluorescence) or astroglia marker GFAP (bottom panel with green fluorescence) although contacts between microglial cells or astrocytes and these axons were observed. The enlarged axons were indicated with yellow arrows and were clearly identifiable with phase contrast microscopy. Scale bar, 50 μm.

**Supplementary Figure 5.**
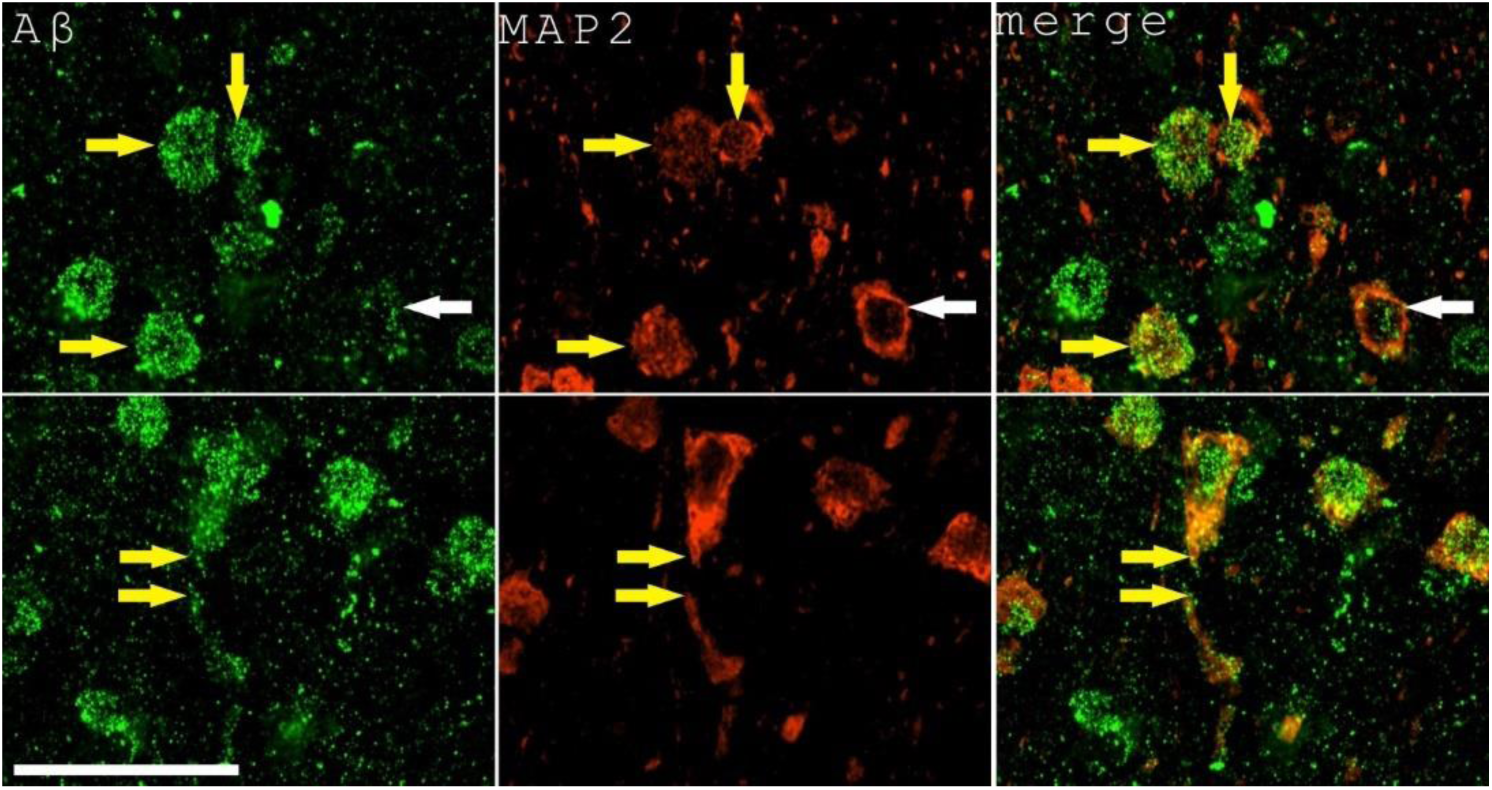
Intracellular Aβ associates with the decrease of MAP2 expression in neurons in AD frontal brain tissues. Aβ was labeled with green fluorescence while MAP2 was labeled with red fluorescence. The top panel indicates several neurons (indicated with yellow arrows) with abundant intracellular Aβ had reduced MAP2 staining comparing to a control neuron with little intracellular Aβ (indicated with a white arrow). The bottom panel showed an abrupt loss of MAP2 expression in a proximal axon neighboring region in a large pyramid neuron (indicated with yellow arrows). Scale bar, 50 μm.

**Supplementary Figure 6.**
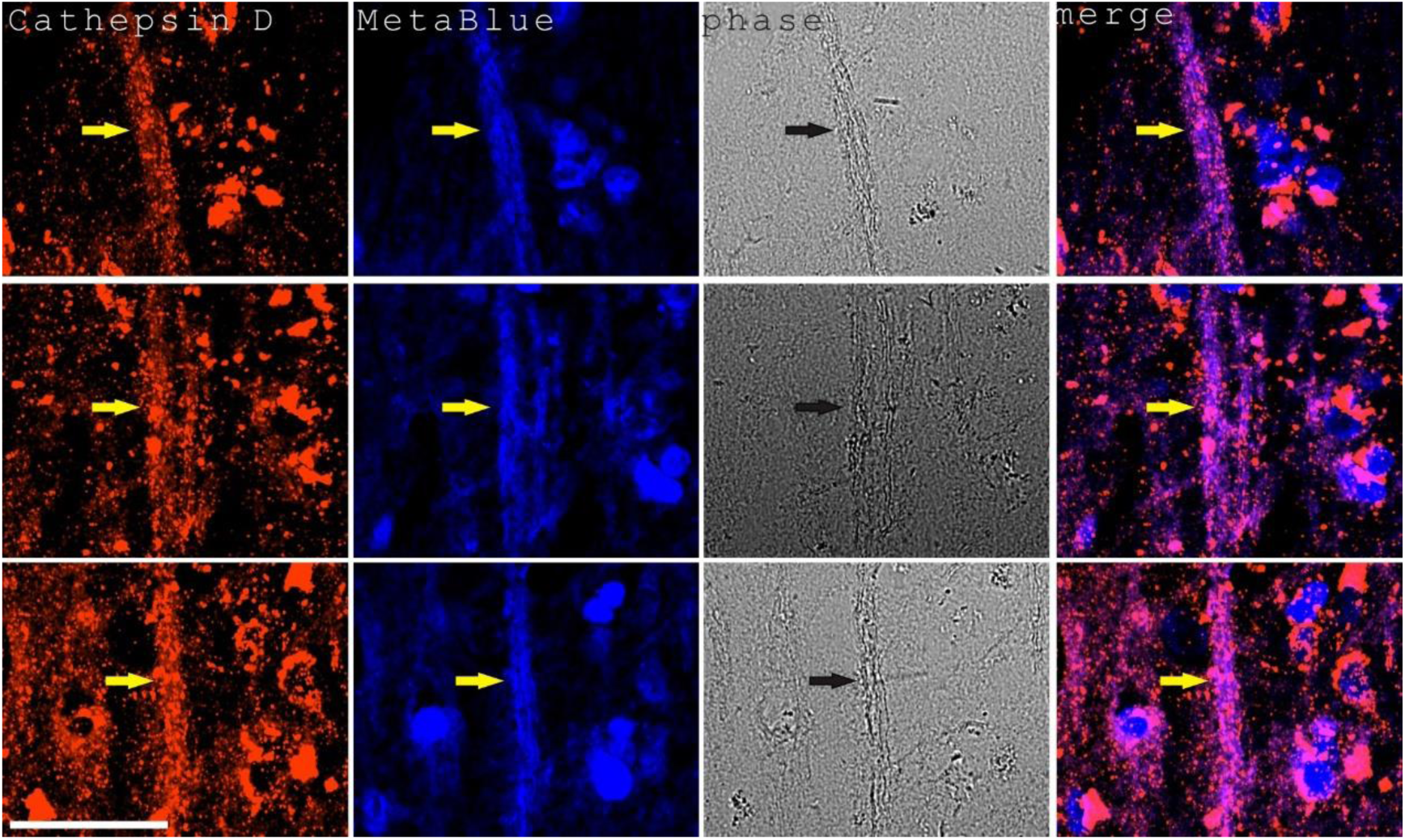
Axon amyloidosis associated with fragmented, very diffusive staining of lysosome marker Cathepsin D in AD frontal brain tissues. Cathepsin D was labeled with red fluorescence while amyloid blue autofluorescence (MetaBlue) was used as a marker for axon amyloidosis. This section was also stained with DAPI to show the nuclear staining. Three representative images were shown. The arrows indicated axon bundles with amyloidosis and very diffusive Cathepsin D staining. Scale bar, 50 μm.

**Supplementary Figure 7.**
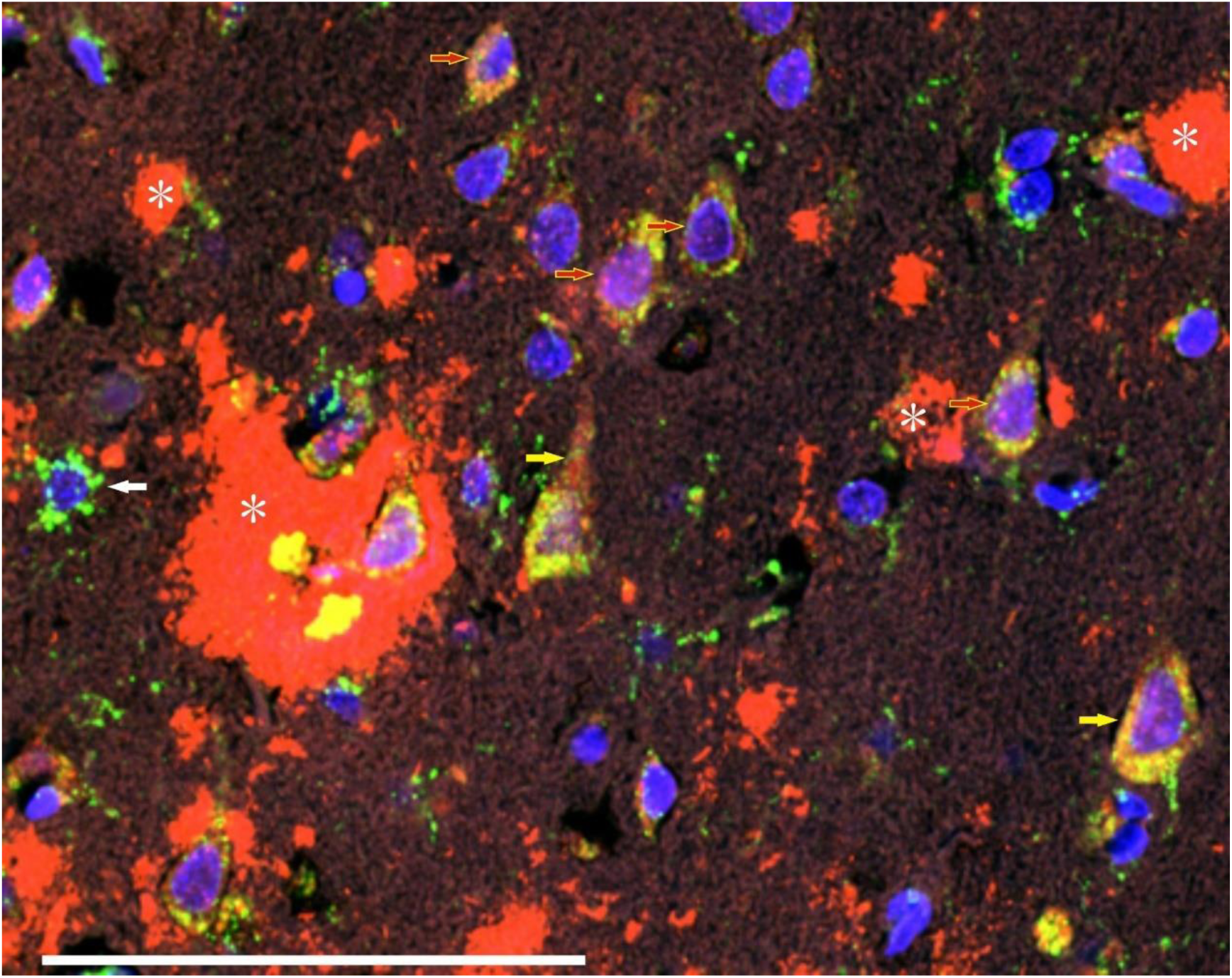
A representative overview image of Aβ and Cathepsin D expression in neural cells in AD frontal brain tissues. Aβ was stained in red fluorescence while Cathepsin D was stained in green. The nuclei was labeled in blue with DAPI. Two large pyramid neurons were indicated with yellow arrows, with the left one showing abundant Aβ staining in the somatic region extending to the proximal axon. A likely astral glial cell was indicated with the white arrow as the Cathepsin D staining indicated the outspreading astral processes. Several senile plaques were indicated with asterisks. Several other type neural cells were indicated with red arrows showing Aβ and Cathepsin D staining signals. Scale bar, 100 μm.

**Supplementary Figure 8.**
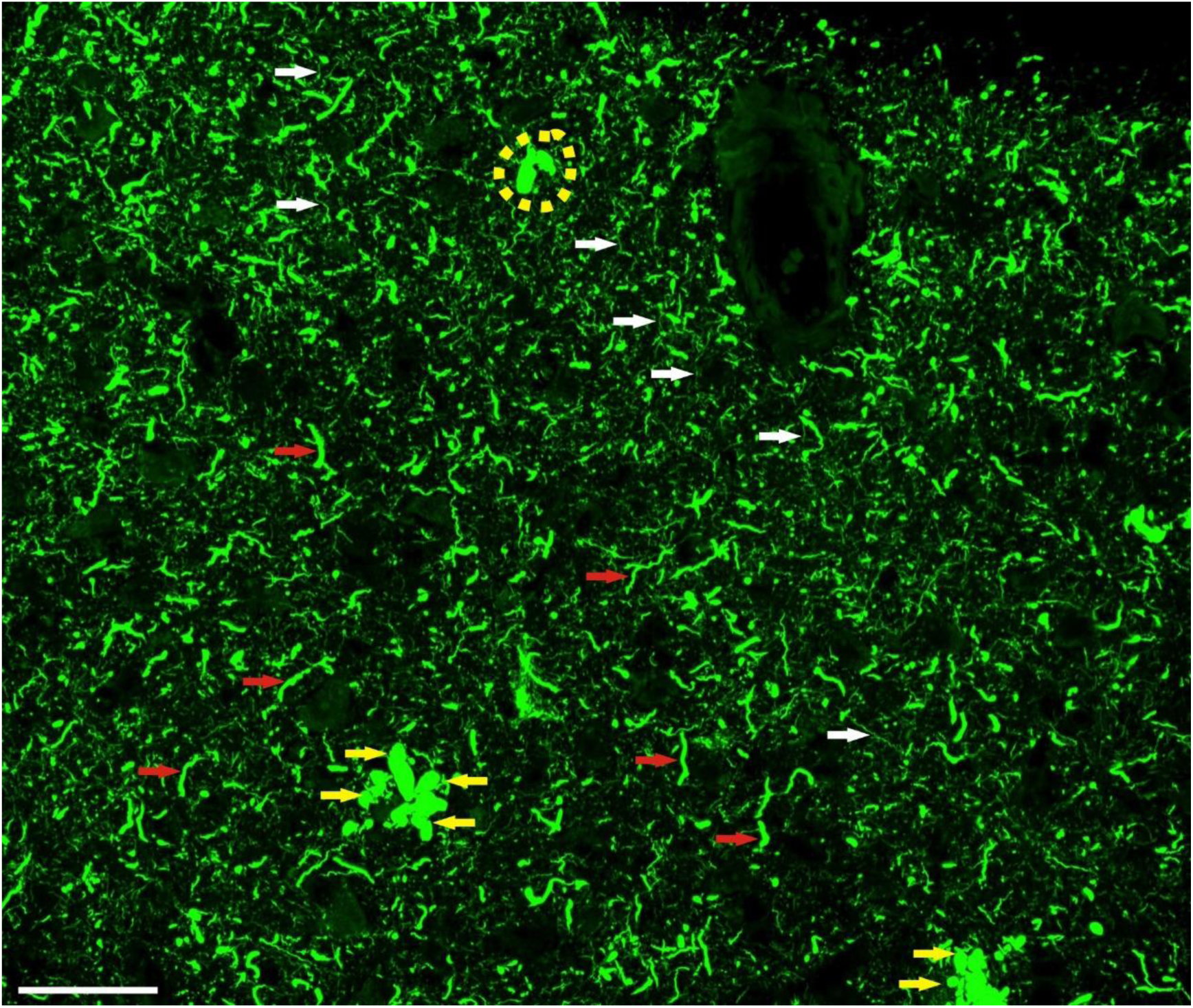
Tau phosphorylation antibody stained abundant dystrophic neurites widely distributed in AD frontal brain tissues, not restricted to dystrophic neurites surrounding the senile plaques and tangled neurons. Senile plaque dystrophic neurites labeled with green fluorescence by phos-Tau antibody were indicated with yellow arrows. A tangled neuron was indicated with a yellow dashed circle. Widely distributed dystrophic neurites were indicated with red arrows. Secondary, much smaller dystrophic neurites were indicated with white arrows. Primary dystrophic neurites had an average diameter of 1.56±0.37 μm (N=40) and secondary dystrophic neurites had an average diameter of 0.40±0.09 μm (N=30). Scale bar, 50 μm.

**Supplementary Figure 9.**
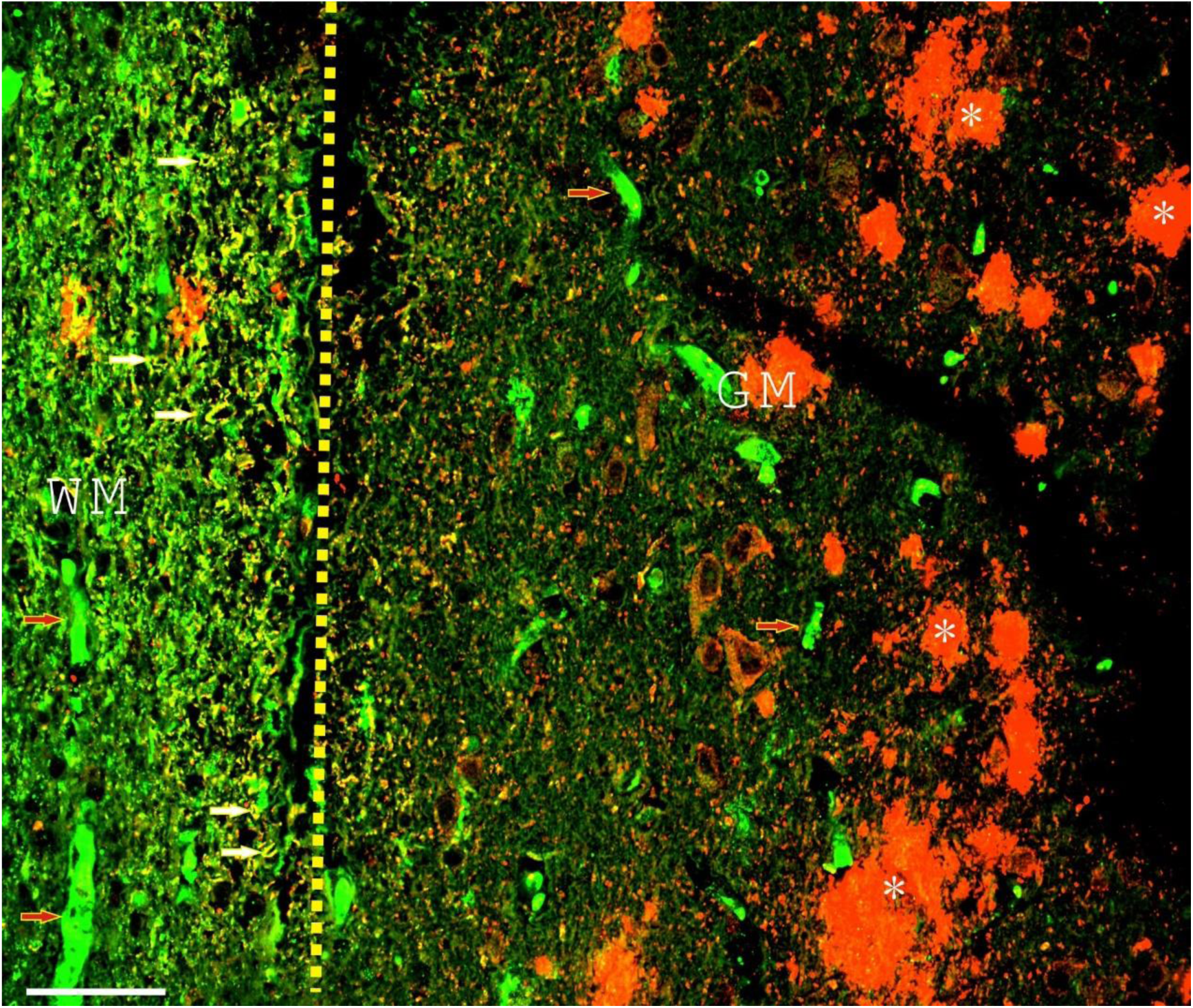
An overview image of HBA and Aβ staining in AD frontal brain sections showing abundant HBA accumulation in the brain white matter axons (HBA labeled green while Aβ labeled red). Yellow dashed lines showed the separation between grey matter (on the right) and white matter (on the left). Blood vessels in white matter and grey matter were all clearly labeled with red blood cell marker HBA (indicated with red arrows). At the same time, strong signals in the white matter axons were observed (some representative axons were indicated with white arrows). Senile plaques were marked with asterisks. WM, white matter. GM, grey matter. Scale bar, 50 µm.

**Supplementary Figure 10:**
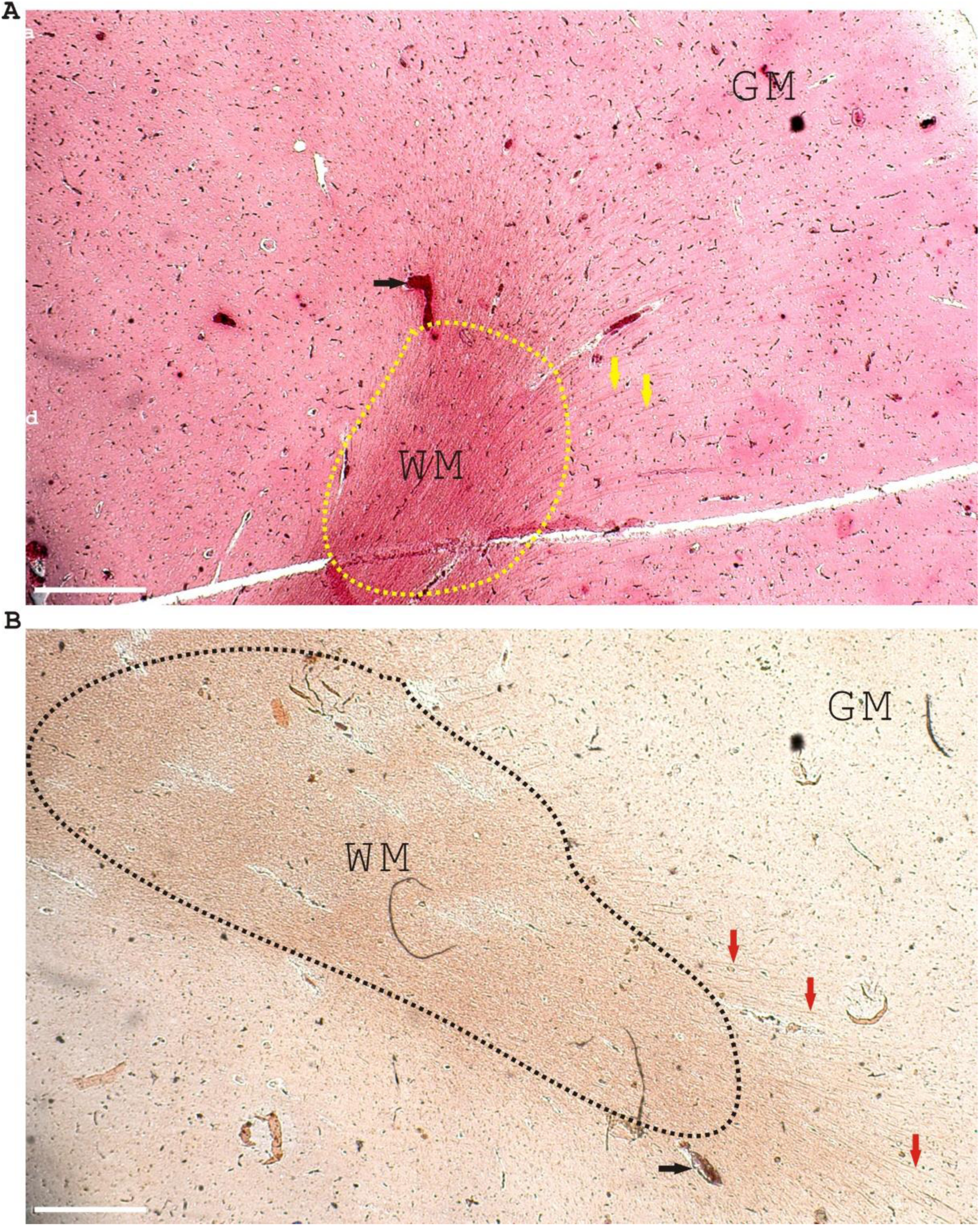
Many axons were positively stained by Alizarin Red and Rhodanine in AD frontal brain tissues. **A.** This panel showed that Alizarin red positively stained the blood cells in the blood vessels (a representative blood vessel was indicated with a black arrow) and the axons (indicated with yellow arrows). Alizarin red stained the white matter (WM) (indicated with yellow dashed circles) clearly while there was less staining in the grey matter (GM). **B.** This panel showed that Rhodanine positively stained the blood cells in the blood vessels (a representative blood vessel was indicated with a black arrow) and the axons (indicated with red arrows). The white matter was also labeled with Rhodanine broadly (indicated with black dashed circles). GM, grey matter. WM, white matter. Scale bar, 500 μm.

**Supplementary Figure 11.**
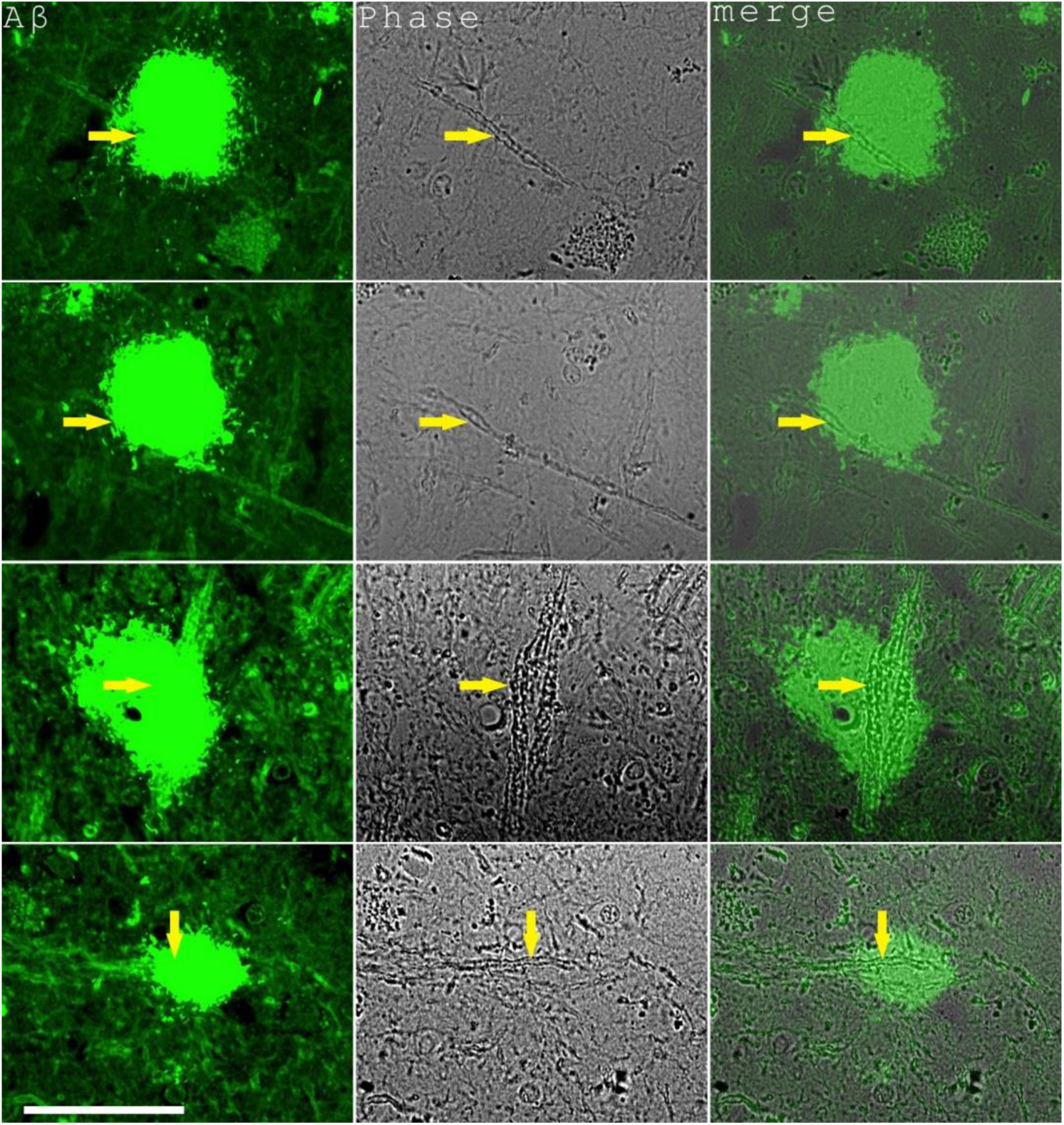
Enlarged axons (indicated with arrows) were detected inside of senile plaques in AD frontal brain tissues but axon breakage was not readily observed. Four representative images were shown. Scale bar, 50 μm.

## Notes

### Competing Interest Statement

The authors have declared no competing interest.

### Summary of Updates

Figures revised. References updated. Manuscript text revised. Supplementary files updated.

